# Repeated parallel losses of inflexed stamens in Moraceae: phylogenomics and generic revision of the tribe Moreae and the reinstatement of the tribe Olmedieae (Moraceae)

**DOI:** 10.1101/2020.04.08.030452

**Authors:** Elliot M. Gardner, Mira Garner, Robyn Cowan, Steven Dodsworth, Niroshini Epitawalage, Deby Arifiani, S. Sahromi, William J. Baker, Felix Forest, Olivier Maurin, Nyree J.C. Zerega, Alexandre Monro, Andrew L. Hipp

## Abstract

We present a densely-sampled phylogenomic study of the mulberry tribe (Moreae, Moraceae), an economically important clade with a global distribution, revealing multiple losses of inflexed stamens, a character traditionally used to circumscribe Moreae. Inflexed stamens facilitate ballistic pollen release and are associated with wind pollination, and the results presented here suggest that losses of this character state may have evolved repeatedly in Moraceae. Neither Moreae nor several of its major genera (*Morus*, *Streblus*, *Trophis*) were found to be monophyletic. A revised system for a monophyletic Moreae is presented, including the reinstatement of the genera *Ampalis, Maillardia, Taxotrophis,* and *Paratrophis*, and the recognition of the new genus *Afromorus*, based on *Morus* subgenus *Afromorus*. *Pseudostreblus* is reinstated and transferred to the Parartocarpeae, and *Sloetiopsis* is reinstated and transferred to the Dorstenieae. The tribe Olmediae is reinstated, replacing the Castilleae, owing to the reinstatement of the type genus *Olmedia,* and its exclusion from Moreae. *Streblus* s.s. is excluded from Moreae and transferred to the Olmediae, which is characterized primarily by involucrate inflorescences without regard to stamen position. Eight new combinations are made.

## Introduction

The preservation of plesiomorphic (ancestral) characters can result in species that are similar in appearance but distantly related, connected only by a remote common ancestor. The mulberry family (Moraceae Gaudich., seven tribes, ca. 39 genera and 1,200 species) illustrates this principle well. Inflexed stamens in bud—an adaptation to wind pollination that allows explosive pollen dispersal when flowers open—were traditionally used to define a tribe of the family, the Moreae (mulberries and their allies). Yet phylogenetic analyses have revealed that inflexed stamens, an ancestral feature of both Moraceae and their sister family Urticaceae (nettles), have been lost repeatedly (Clement & Weiblen, 2009). Thus, for example, Cecropiaceae C.C. Berg, traditionally distinguished from the nettle family by the absence of inflexed stamens, is in fact embedded within the Urticaceae Juss. (Berg, 1978; Clement & Weiblen, 2009). Likewise, while mulberries (*Morus* L.), paper mulberries (*Broussonetia* L’Hér. ex Vent.), and osage oranges (*Maclura* Nutt.) were once treated as tribe Moreae Gaudich. on account of their inflexed stamens, phylogenetic analyses reveal them to belong to three distinct clades, each containing an assemblage of genera with and without inflexed stamens (Clement & Weiblen, 2009; Zerega & Gardner, 2019).

This study focuses on the tribe Moreae, a clade of six genera and an estimated 66 species (Clement & Weiblen, 2009). This widespread group of plants contains species of ecological and cultural importance, as well as the economically important white mulberry (*Morus alba* L.), whose leaves sustain *Bombyx mori* L. (Bombycidae), the invertebrate proletariat of the silk industry.

### Morphological basis for higher classification in Moraceae

By the end of the nineteenth century, Engler (1889) had circumscribed a Moraceae that is quite similar to the modern concept of the family, with two subfamilies: the Moroideae, comprising tribes Dorstenieae Gaudich., Broussonetieae Gaudich., Fatouae Engler, Moreae, and Strebleae Bureau, characterized by stamens inflexed in bud (broadly construed, including *Dorstenia* where they straighten gradually rather than spring outward suddenly); and the Artocarpoideae, composed of tribes Brosimae Trécul, Euartocarpeae Trécul, Ficeae Dumort., and Olmedieae Trécul and characterized by stamens straight in bud. The two most influential scholars of Moraceae classification in the 20th century were E.J.H. Corner and C.C. Berg. Corner considered inflorescence architecture to be the most important character for higher-rank taxonomy within the family and inflexed stamens consequently of secondary importance(Corner, 1962). Berg by contrast questioned the utility of inflorescence architecture and took into account a variety of characters, especially the presence of inflexed stamens (Berg, 1977b, 2001).

The taxonomic history of the family has leaned heavily on this stamen character. Engler’s Moreae (1899), unchanged in substance from Bureau’s (1873), contained six genera, *Ampalis* Bojer*, Pachytrophe* Bureau*, Paratrophis* Blume*, Pseudomoru*s Bureau, and *Morus*, all with inflexed stamens that spring out suddenly at anthesis (Table 1). Corner’s expanded Moreae consisted of seven genera with either straight or inflexed stamens, but with pistillate inflorescences never condensed into a head: *Fatoua* Gaudich, *Morus*, *Sorocea* A. St.-Hil. (apparently including *Paraclarisia* Ducke), *Clarisia* Ruiz & Pav., *Ampalis*, *Pachytrophe*, and *Streblus* Lour. (including *Bleekrodea* Blume*, Paratrophis, Pseudomorus*, *Taxotrophis* Blume, *Sloetiopsis* Engl. and *Neosloetiopsis* Engl.). Corner himself, however, found the diversity of his Moreae unsatisfactory, noting that “[t]oo many genera on insufficient and invalid grounds trouble this small tribe” (Corner, 1962). Perhaps in response, Berg’s Moreae comprised all of the genera with inflexed stamens and none without, including *Broussonetia* and *Maclura* but excluding *Sorocea* and *Clarisia*, providing a simple character with which to delimit the tribe(Berg, 2001; Berg & al., 2006).

**Table 1.**
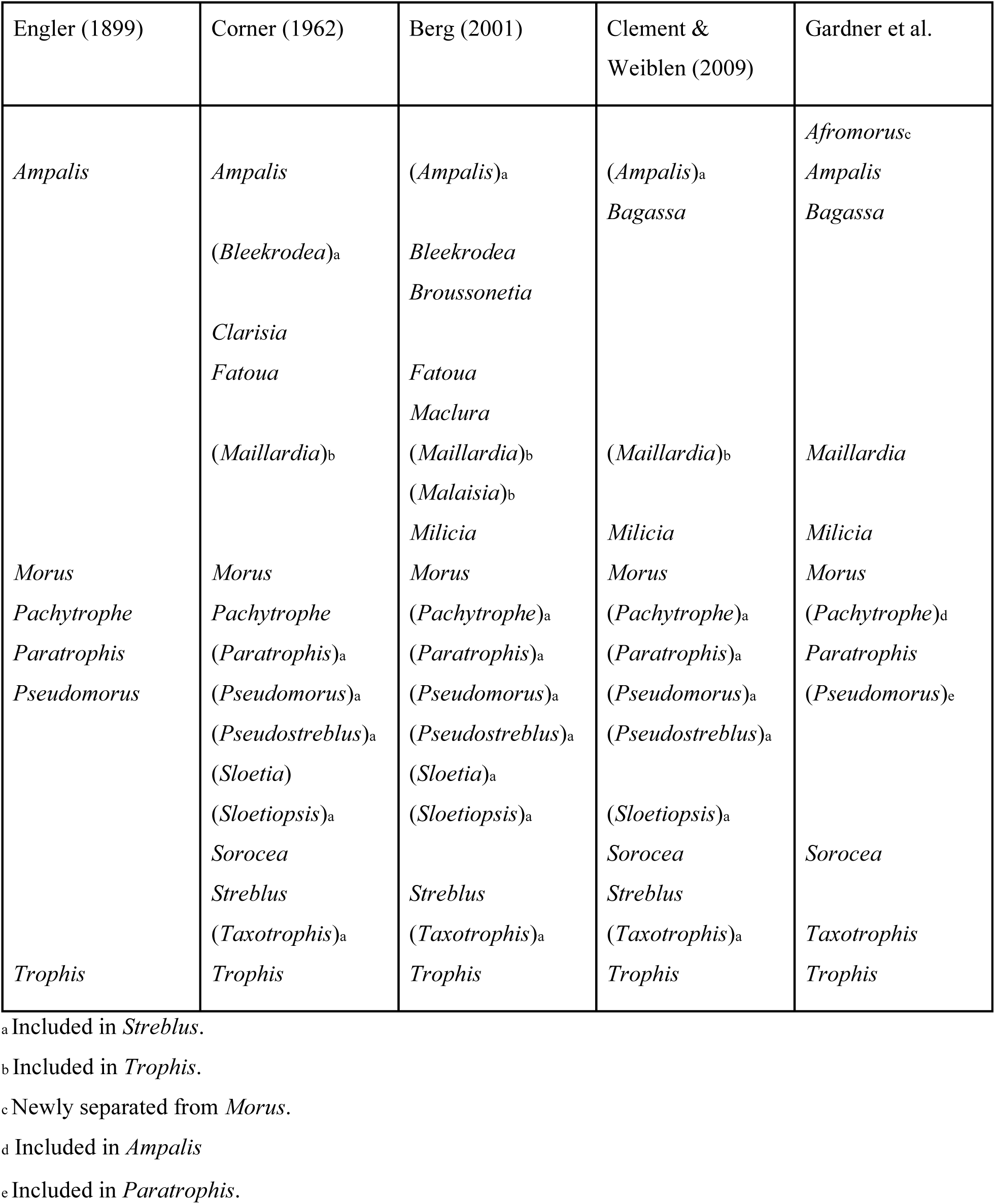
Overview of the taxonomic history of Moreae.

Recent phylogenetic work has supported a third approach, with the Moreae comprising six genera, including genera with both straight (*Sorocea*, *Bagassa* Aubl.) and inflexed (*Broussonetia*, *Maclura*) stamens (Clement & Weiblen, 2009). These studies also suggest that the character state of inflexed stamens is plesiomorphic and is the ancestral state for both Moraceae and Urticaceae (Datwyler & Weiblen, 2004; Zerega & al., 2005; Clement & Weiblen, 2009). Inflexed stamens, which are precursors to ballistic pollen release, are associated with wind pollination (Bawa & Crisp, 1980; Berg, 2001), while the loss of inflexed stamens is associated with animal pollination. Although dominant in the Moreae, inflexed stamens also occur in two other tribes: Maclureae W.L. Clement & G. Weiblen and Dorstenieae. Genera lacking inflexed stamens occur in all seven tribes of Moraceae (Clement & Weiblen, 2009; Zerega & Gardner, 2019).

### Taxonomic summary of Moreae

Following Clement & Weiblen (2009), the Moreae comprise six genera and an estimated 66 species: *Bagassa* (1), *Milicia* Sim (2), *Morus* (16), *Sorocea* (16), *Streblus* (23), and *Trophis* P. Browne. (8) (Berg, 1977a, 2001; Berg & al., 2006; Clement & Weiblen, 2009; Filho & al., 2009; Machado & al., 2013; Santos & Neto, 2015). Here, we present an overview of these genera.

*Morus* L., the true mulberries, is characterized by leaves with crenate margins and trinerved bases, stamens inflexed in bud, and many-flowered pistillate spikes whose four-parted perianths become fleshy in fruit, the aggregations superficially resembling a blackberry. *Morus* comprises approximately 16 species whose delimitation requires further research. There are three subgenera: *Morus* (ca. 14 species) which is found in temperate to tropical Asia and from North America to Mexico; *Gomphomorus* Leroy, a single species restricted to tropical South America; and *Afromorus* (Bureau ex Leroy), a single species restricted to tropical Africa. Previous phylogenetic work has suggested that these three subgenera may not form a monophyletic clade (Nepal, 2012). *Milicia* (2 spp., Africa) has inflorescences which somewhat resemble those of *Morus*, but the leaves of *Milicia*, with entire margins and pinnate venation, prevent any confusion of the two genera.

*Streblus* Lour., with 23 species from India to Southeast Asia and Oceania, is morphologically heterogeneous, but its species are all characterized by stamens inflexed in bud and pistillate flowers with more or less free tepals that enclose the fruit loosely or not at all. Initially described by Loureiro based on the widespread *S. asper* Lour.—notable for its discoid-capitate staminate inflorescences with the rudiments of an involucre—*Streblus* was broadened by Corner (1962), bringing in as sections *Taxotrophis*, *Phyllochlamys* Bureau, *Paratrophis* (including *Pseudomorus*), *Pseudostreblus* Bureau, *Bleekrodea*, and *Sloetia* Teijsm. & Binn. (apparently including *Sloetiopsis* but without making any combinations). None of these have discoid-capitate inflorescences; they mostly have spicate staminate inflorescences, and the latter two can have bisexual inflorescences. Corner viewed these sections as fragments of an ancient lineage preserving ancestral characters (Corner, 1975). Following additional work by Corner (1970, 1975), Berg included the genera *Ampalis* and *Pachytrophe* as section *Ampalis* (Bojer) C.C. Berg, subsumed *Sloetiopsis* and section *Taxotrophis* into section *Streblus*, and excluded section *Bleekrodea*, reinstating it as a genus (Berg, 1988). Berg later reinstated section *Taxotrophis*, whose species are unique in bearing axillary spines (Berg, 2005; Berg & al., 2006). In 2009, Clement and Weiblen reinstated *Sloetia* at generic rank and transferred it and *Bleekrodea* to Dorstenieae based on phylogenetic evidence (Clement & Weiblen, 2009).

*Trophis* P. Browne is characterized by stamens inflexed in bud, spicate staminate inflorescences, and tubular pistillate perianths, becoming fleshy and enclosing the fruit, except for the monotypic section *Olmedia* (Ruiz & Pav.) C.C. Berg (*T. caucana*), which has discoid-capitate staminate inflorescences with an involucre. *Trophis* (regrettably conserved over Linnaeus’s *Bucephalon* L.) as recognized by Berg (1988, 2001) had five sections: *Trophis*, restricted to Latin America; *Calpidochlamys* (Diels) Corner (previously included in *Paratrophis* and *Uromorus*), restricted to Southeast Asia (Corner, 1962); *Maillardia* (Frapp. ex Duch.) C.C. Berg, restricted to Africa; *Malaisia* (Blanco) C.C. Berg, restricted to Southeast Asia; and *Olmedia*, restricted to Latin America. The sinking of *Olmedia* into *Trophis* by Berg (1988), requiring the re-typification of the tribe Olmedieae Trécul, which became Castilleae C.C. Berg. Recently, *Malaisia* Blanco was reinstated as a genus and transferred to Dorstenieae based on phylogenetic evidence (Clement & Weiblen, 2009), reducing the current number of sections to four.

Two Neotropical genera included in Moreae by Clement and Weiblen (2009) but not by Berg (2001) have straight stamens. While the monotypic *Bagassa* Aubl., with its long staminate catkins, is wind pollinated (Bawa & Crisp, 1980), evidence suggests that *Sorocea* A. St.-Hil. (19 spp.), which produces racemose staminate inflorescences that do not have as many flowers as those of *Bagassa*, likely contains both wind- and insect-pollinated species (Zapata & Arroyo, 1978; Bawa & al., 1985; Lewis, 1986). The pistillate perianths are subtended by pluricellular hairs, believed to serve as a substrate for a fungus which, in turn attracts pollinators (Berg, 2001). The staminate flowers of *Sorocea affinis* Helmsl. are apparently fragrant (fide *S. Zona 778*, 21 Nov. 1997, FTBG♂♀), suggesting insect pollination for that species. If insect pollination is confirmed in *Sorocea*, it would likely represent another transition from ancestral wind to derived insect pollination in Moraceae.

Disagreement between the two principal recent monographers of the Moraceae, Corner (1962) and Berg (2001, 2005, 2006), over the delimitation of the Moreae has been compounded by a series of phylogenies based on the analysis of molecular (DNA sequence) data that have redefined genera and their relationships. This has resulted in a poorly delimited tribe, both with respect to diagnostic morphological characters, the genera it comprises, and the rank of several taxa (see above). Ensuring that the tribe and its genera are monophyletic will result in a classification that better reflects evolutionary history, providing a framework for answering broader scientific questions. Our aim, therefore, was to generate a comprehensive phylogeny of the Moreae (Fig. 1) and its near allies by sampling all of the potential genera and species in the tribe sensu Berg, and sensu Clement & Weiblen (2009) and use this to redelimit the tribe and its genera. We aimed to do so through increased taxon and genome sampling compared to previous studies, including a nearly comprehensive sample of the taxa in Moreae (56/66 species) and the allied tribes Artocarpeae (76/83 taxa) and Maclureae (12/12). Our data set combines phylogenomic data generated using two largely non-overlapping sets of enrichment baits, one developed specifically for the Moraceae (Gardner et al., 2016), the other for the whole of the Angiosperms (Johnson & al., 2019), allowing us to explore the possibilities and challenges of combining samples based on largely non-overlapping loci. The resulting generic revision lays the groundwork for species-level revisionary work and provides clarity to this economically important clade.

**Figure 1.**
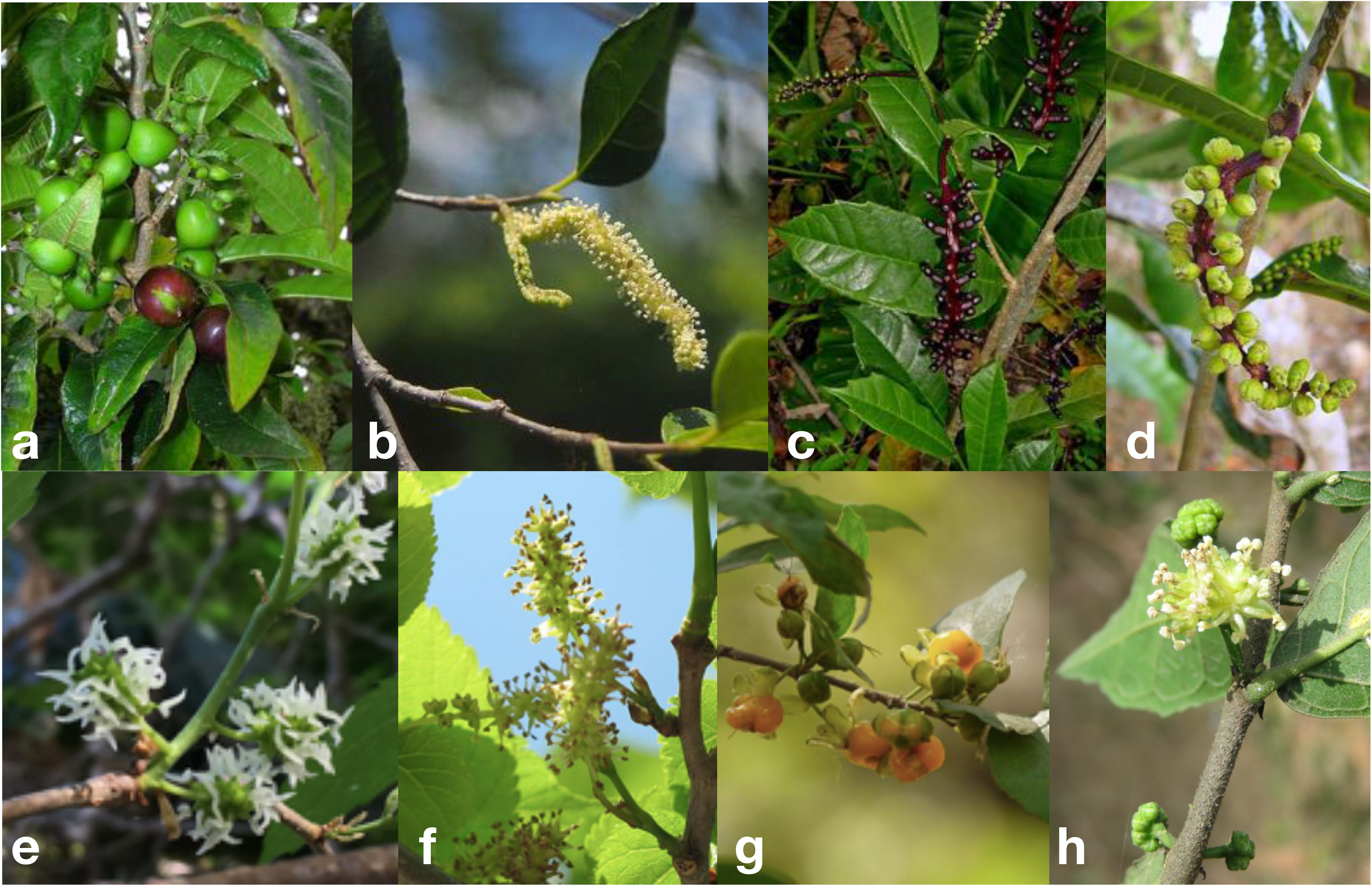
Representatives of Moreae: *Paratrophis pendulina* (=*Streblus pendulinus*) (a) infructescences and (b) staminate inflorescences, the latter showing the characteristic sterile groove; *Sorocea racemosa* (c) infructescences and (d) staminate inflorescence; *Morus nigra* (e) pistillate inflorescences with elaborate stigmas and (f) staminate inflorescences with stamens that, having been inflexed in bud, are substantially longer than the perianth itself; and a newly-placed member of Olmedieae (=Castilleae), *Streblus asper:* (g) infructescences and (h) discoid-capitate staminate inflorescences with the rudiments of an involucre visible below the unopened flowers. Photo credits: (a) F. & Kim Starr, under a CC-BY-3.0-US licence (https://commons.wikimedia.org/wiki/File:Starr-051029-5102-Streblus_pendulinus-fruit-Auwahi-Maui_(24481336159).jpg); (b) M. Marathon, CC-BY-SA-4.0 (https://commons.wikimedia.org/wiki/File:Streblus_brunonianus_flowers.jpg); (c)-(d) A. Popovkin, CC-BY-2.0 (https://commons.wikimedia.org/wiki/File:Sorocea_racemosa_Gaudich._-_Flickr_-_Alex_Popovkin,_Bahia,_Brazil_(3).jpg and https://commons.wikimedia.org/wiki/File:Sorocea_racemosa_Gaudich._-_Flickr_-_Alex_Popovkin,_Bahia,_Brazil_(9).jpg); (e) E. Gardner; (f) Schurdl, CC-BY-SA-4.0 (https://commons.wikimedia.org/wiki/File:Morus_nigra_100525_02.jpg); (g) D. Valke, CC-BY-SA-2.0 (https://commons.wikimedia.org/wiki/File:Bekar_(Konkani-_बेकर)_(4533519363).jpg); (h) Vinayaraj, CC BY-SA 4.0 (https://commons.wikimedia.org/wiki/File:Streblus_asper_at_Panamaram_(5).jpg)

We also set out to reconstruct the evolutionary history of inflexed stamens within Moraceae using ancestral state reconstruction. The loss of inflexed stamens in Castilleae+Ficeae and Artocarpeae have been associated with transitions from wind to animal pollination (Momose & al., 1998; Sakai & al., 2000; Datwyler & Weiblen, 2004; Gardner & al., 2018). A more complete picture of evolutionary transitions between inflexed and straight stamens may help focus further research on transitions in pollination biology with Moraceae.

## Materials and Methods

We used target enrichment sequencing (HybSeq) (Weitemier & al., 2014) to capture 333 genes previously developed for phylogenetic work in Moraceae (the “Moraceae333”) (Gardner & al., 2016; Johnson & al., 2016). This method allows for efficient capture of hundreds of loci and is suitable for both fresh material and degraded DNA from herbarium material (Villaverde & al., 2018; Brewer & al., 2019), which comprises much of the material employed in this study.

### Taxon sampling, library preparation, and sequencing

We sampled 56 out of 66 species (and all genera) in Moreae as well as select outgroup taxa from Artocarpeae using DNA from leaf tissue preserved on silica gel or—in most cases—samples from herbarium specimens (Table S1). In either case, DNA was extracted using a modified CTAB method, usually with increased incubation times to maximize yield from herbarium tissue(Doyle & Doyle, 1987; Hale & al., 2020). Samples were quantified using a Qubit fluorometer (Invitrogen, Life Technologies, California, USA), and herbarium samples were run on a gel to test for degradation. For most samples 200 ng of DNA was used for library preparation; for some low-yield samples, as little as 50 ng was used, and for very degraded samples, as much input as possible was used, up to 400 ng. Undegraded DNA was fragmented either using NEB DNA Fragmentase (New England Biolabs, Ipswich, Massachusetts, USA) or on a Covaris M220 (Covaris, Wobum, Massachusetts, USA). DNA samples with an average fragment size less than 500 bp were not fragmented at all, and partially-degraded samples with an average fragment size of over 500 bp were fragmented on the Covaris M220. TruSeq-style library preparation was carried out using either the KAPA Hyper Prep kit (Kapa Biosystems, Wilmington, MA) or the NEB DNA Ultra 2 kit following the manufacturer’s protocols, except that end repair, A-tailing, and adapter ligation were carried out in reduced-volume reactions (0.25x for KAPA and 0.5x for NEB) to reduce costs. Final products were quantified on the Qubit and combined in equal molecular weights into pools of 16–20 samples. The pools totaled 1,200 µg each if enough library preparation was available. Pools were hybridized for 16–24 hours to custom Moraceae probes (Gardner et al, 2016) using a MYBaits kit (Arbor Biosciences, Ann Arbor, Michigan, USA) following the manufacturer’s protocol, except that the probes were diluted 1:1 with nuclease-free water. Hybridization products were reamplified using KAPA Hot Start PCR reagents following the MYbaits protocol, quantified on the Qubit, and quality-checked on an Agilent BioAnalyzer (Agilent Technologies, Palo Alto, California, USA). Samples with adapter dimer peaks were cleaned using 0.7x SPRI beads and re-run on the Qubit and BioAnalyzer. An initial sequencing run took place on a MiSeq 2 × 300bp run (v3) (Illumina, San Diego, California, USA) at the Field Museum of Natural History, and then additional samples were sequenced on a HiSeq 4000 2 x 150bp run at the Northwestern University Genomics Core.

We also included 47 Moraceae and Urticaceae samples enriched for the Angiosperm353 probes and sequenced as part of the Plant and Fungal Trees of Life project (PAFTOL, RBG Kew; https://www.kew.org/science/our-science/projects/plant-and-fungal-trees-of-life; Table S1). Sample preparation and sequencing followed Johnson et al. (2019). The PAFTOL samples were enriched with a universal probe set developed for angiosperms (the “Angiosperms353”).

Finally, we used samples sequenced for previous phylogenetics projects in *Artocarpus* J.R. Forst. & G. Forst. and Parartocarpeae Zerega & E.M. Gardner to complete our sampling (Johnson & al., 2016; Gardner, 2017; Kates & al., 2018; Zerega & Gardner, 2019). The final dataset contained 247 samples.

### Assembly of reads

We trimmed reads using Trimmomatic (ILLUMINACLIP: TruSeq3-PE.fa:2:30:10 HEADCROP:3 LEADING:30 TRAILING:25 SLIDINGWINDOW:4:25 MINLEN:20) (Bolger & al., 2014) and assembled them with HybPiper, which produces gene-by-gene, reference-guided, *de novo* assemblies (Johnson & al., 2016). For the samples enriched with the Angiosperms353 baits, we used the reference described by Johnson et al. (2019), and for the samples enriched with the Moraceae333 baits, we used the reference described in Zerega & Gardner (2019). To increase overlap between the two data sets beyond the 5 genes they inherently have in common, we used HybPiper to assemble the Angiosperms353-enriched reads using the Moraceae333 targets and vice-versa, in order to capture any additional genes found in off-target reads. The exception was most of the *Artocarpus* data set, which was previously assembled for another study using only the Moraceae333 targets. We used the HybPiper script “intronerate.py” to build “supercontig” sequences for each gene, consisting of exons as well as any assembled flanking non-coding sequences (intronic or intergenic). To these new assemblies, we added 76 Moraceae333 assemblies from Gardner (2017) to complete sampling for the Artocarpeae.

### Main phylogenetic analyses

The full data set was analyzed using the “exon” sequences to ensure good alignment across the entire family. To maximize phylogenetic resolution within Moreae, a smaller data set consisting only of Moreae taxa and a single outgroup taxon (*Artocarpus heterophyllus*) was analyzed using the “supercontig” sequences but following the same methodology. For each of the 686 genes assembled, we discarded sequences shorter than 25% of the average length for the gene and discarded genes containing sequences for less than 30 samples. We aligned each gene with MAFFT (Katoh & Standley, 2013)and used Trimal (Capella-Gutiérrez & al., 2009) to discard sequences with an average pairwise identity of less than 0.5 to all other sequences in the alignment (indicative of poor a poor quality sequence that could not be properly aligned) as well as columns containing more than 75% gaps. We used RAxML 8.2.4 (Stamatakis, 2014) to estimate a maximum-likelihood tree for a partitioned supermatrix of all genes as well as for each gene individually (GTRCAT model, 200 bootstrap replicates). We then used ASTRAL-III (Zhang & al., 2017), a summary-coalescent method, to estimate a species tree from all gene trees, estimating node support using both bootstrap (160 replicates, resampling across and within genes) and local posterior probability (normalized quartet scores, representing gene tree concordance). Finally, we used ASTRAL-III to test whether, for each node, the null hypothesis of a polytomy could be rejected (Sayyari & Mirarab, 2018). Alignment, trimming, and estimation of gene trees were parallelized using GNU Parallel (Tange, 2018).

### Whole chloroplast phylogenetic tree

We also built a whole-chloroplast phylogenetic tree as follows. Rather than assembling full-length genomes, which can be extremely slow to align, we assembled and aligned the genomes in sections, dramatically speeding up the process. We created HybPiper targets using the chloroplast genome of *Morus indica* L. (NCBI RefSeq accession no. NC_008359.1) and the associated gene annotations. Each target consisted of a gene feature concatenated with any internal or subsequent non-coding sequence, terminating one base before the next gene feature began. These intervals were generated by manually editing the NCBI gff3 file in Excel (Microsoft Corp., Redmond, Washington, USA) and then extracted using BedTools (Quinlan & Hall, 2010). Assemblies for all Moraceae samples and *Boehmeria nivea* (L.) Gaudich. were carried out in HybPiper using a coverage cutoff of 2 and otherwise with default parameters. Within each target, sequencing less than 25% of the average length were discarded, and after alignment with MAFFT, sequences with an average pairwise identity of less than 0.7 were discarded (the higher cutoff reflecting the conserved nature of chloroplast DNA). Alignments were then concatenated into a supermatrix, and samples with more than 50% undetermined characters were discarded. Alignments were then rebuilt, filtered, and concatenated, and columns in the final supermatrix with more than 75% missing characters were discarded. A maximum-likelihood tree was generated using RAxML 8.2.4 under the GTRCAT model, with 200 rapid bootstrap replicates.

### Phylogenetic network analysis

To further investigate relationships within the *Paratrophis* clade, including the proper placement of *Morus insignis* and *Trophis philippinensis* (Bureau) Corner, we constructed phylogenetic networks based on two reduced 8-taxon datasets (“exon” and “supercontig”) consisting of those taxa plus *Streblus anthropophagorum* (Seem.) Corner*, S. glaber* (Merr.) Corner*, S. glaber* subsp. *australianus* (C.T. White) C.C. Berg, and *S. heterophyllus* (Blume) Corner, with *S. mauritianus* (Jacq.) Blume as the outgroup. Alignment preparation followed the workflow outline above except that only genes with sequences for all eight taxa were retained. Rooted gene trees were used to infer the network in PhyloNet using the “InferNetwork_MPL” command, allowing a maximum of four hybridization events and collapsing gene tree nodes with less than 30% bootstrap support (Yu & Nakhleh, 2015; Wen & al., 2018) (Wen et al., 2018; Yu and Nakhleh, 2015); AIC was scored for the best five networks from 10 runs, using the number of branch lengths and hybridization events calculated as the number of parameters (Kamneva & al., 2017).

### ITS and rbcL phylogenetic trees

While we did not have our own sequences for *Streblus tonkinensis* (Eberh. & Dubard) Corner, *S. ascendens* Corner, *S. banksii* (Cheeseman) C.J. Webb, and *S. smithii* (Cheeseman) Corner, either *ITS* or *rbcL* sequences existed in NCBI GenBank for these taxa (Fig. S2). We were not able to examine the underlying specimens ourselves, but we considered the chances of misidentification low because those species are morphologically and/or geographically distinctive. We constructed *ITS* and *rbcL* data sets from our own samples using HybPiper, using *Streblus* sequences for these loci (obtained from GenBank) as targets and reducing the coverage cutoff at 2 to recover the loci from off-target reads. To these sequences, we added GenBank sequences of *Streblus* taxa as well as *Trophis caucana* (Pittier) C.C. Berg. For each locus, we aligned sequences with MAFFT, discarded short sequences (<490 bp), trimmed alignments to remove columns with over 75% gaps, and built maximum-likelihood trees using RAxML (GTRGAMMA, 1000 rapid bootstraps).

### Divergence time estimation

The supermatrix maximum-likelihood tree was time calibrated using ape v 5.3 (Paradis & Schliep, 2019) in R v 3.5.1 (Team, 2019). First, the tree was pruned of duplicate taxa and non-Moraceae outgroup taxa, and edge lengths (in substitutions per site) were multiplied by the number of sites in the alignment (to convert them to substitutions). The following stem nodes were constrained with minimum ages (in million years, Ma) based on fossil data, following(Zhang & al., 2019): *Ficus,* 56 Ma; *Broussonetia*, 33.9 Ma; *Morus* (subg. *Morus*, based on U.S.S.R. locality of the fossil), 33.9 Ma; and *Artocarpus*, 64 Ma. The crown node of Moraceae was constrained to a minimum age of 73.2 Ma and a maximum age of 84.7 Ma.(Zhang & al., 2019), which had a more extensive outgroup sampling that we have here. The tree was then time-calibrated using the chronos function under two different models (“relaxed” and “correlated”) (Kim & Sanderson, 2008; Paradis, 2013). The smoothing parameter (λ) was chosen using the cross-validation method in the chronopl function (testing λ = 0 and 0_-5_ through 10_15_), selecting the value of λ that minimized the cross-validation statistic (Sanderson, 2002). The resulting trees were visualized using the densiTree function in phangorn 2.4.0 (Schliep, 2011), and the time calibrated tree representing the central tendency of these analyses was selected for use in all further analyses, also taking into account the results of past family-wide studies. As the penalized likelihood approach used does not integrate over model uncertainty or uncertainty in calibration placement and timing, confidence intervals on node ages are not provided in this study. A geologic timescale based on the strat2012 dataset added to tree figures using PHYLOCH v 1.5-3 (Heibl, 2008).

### Ancestral state reconstruction for inflexed stamens

All taxa were coded for presence (1) or absence (0) of inflexed stamens in bud. Taxa such as *Dorsteniae* with stamens that are inflexed in bud but gradually straighten were coded as 0. We reconstructed ancestral character states on the entire phylogeny by stochastic mapping using the make.simmap function in the phytools v 0.6-99 (Revell, 2012). We also tested for trait-associated shifts in diversification rates using BAMM v 2.5.0 (Rabosky & al., 2013, 2014a; Rabosky, 2014), specifying the amount of missing taxa per clade to account for the high proportion of missing taxa in Ficeae, Castilleae, and Dorstenieae. Parameters were optimized using BAMMtools v 2.1.7 (Rabosky & al., 2014b). We also tested for trait-dependent diversification using diversitree v 0.9-13 (FitzJohn, 2012). We evaluated BiSSE and trait-independent models on the entire tree and on a pruned tree consisting only of the densely-sampled Maclureae+Moreae+Artocarpeae clade.

Reads from have been deposited in Genbank (Non-PAFTOL samples: BioProject PRJNA322184; PAFTOL samples: BioProject PRJEB35285, subprojects PRJEB37667 for Moraceae and PRJEB37665 for Urticaceae). Tree files and scripts outlining the phylogenetic analyses have been deposited in the Dryad Data Repository (###TBA###).

## Results

The final data set contained 247 samples, 47 enriched with the Angiosperms353 baits, 196 enriched with the Moraceae333 baits, and 4 extracted from whole genomes) and 619 genes (286 Angiosperms353 and all of the Moraceae333) (Table S1). The “exon” supermatrix (all taxa) contained 613,126 characters, and the “supercontig” supermatrix (Moreae and select outgroup taxa only) contained 799,926 characters. The chloroplast data set contained 113 loci.

On average, we assembled 39 Moraceae333 genes from the Angiosperms353-enriched samples and 20 Angiosperms353 genes from the Moraceae333-enriched samples, and the average pairwise overlap in assembled loci was 210 (Tables S1, S2). Nineteen taxa were represented in both sets of samples, and 15 of these were always monophyletic (Figs. 2, 3). Twelve resolved as sister pairs: *Antiaropsis decipiens* (31 loci overlapping), *Bagassa guianensis* Aubl. (38), *Batocarpus amazonicus* (Ducke) Fosberg (22), *Batocarpus orinocensis* H. Karst. (34), *Brosimum alicastrum* Sw. (42), *Clarisia racemosa* Ruiz & Pav. (59), *Maclura africana* (Bureau) Corner (213), *Milicia excelsa* (Welw.) C.C. Berg (45), *Streblus asper* (Retz.) Lour. (75), *Streblus mauritianus* (Jacq.) Blume (40), *Streblus usambarensis* (Engl.) C.C. Berg (43), and *Trophis caucana* (32). Three more represented by more than two samples always resolved as a clade: *Sorocea bonplandii* (Baillon) W.C. Burger, Lanj. & de Boer (3 samples; 19–35 loci overlapping), *Maclura tinctoria* (L.) D. Don ex Steud. (4 samples; 37–43 loci overlapping), and *Malaisia scandens* (Lour.) Planch. (3 samples; 27–43 loci overlapping). Three additional taxa resolved as sister pairs in the ASTRAL analysis but as a grade in the supermatrix analysis: *Maclura cochinchinensis* (Lour.) Corner (28 loci overlapping), *Streblus heterophyllus* (Blume) Corner (50), and *Trophis montana* (Leandri) C.C. Berg (56). Only one species, *Utsetela gabonensis* Pellegr. (18 loci overlapping) was never monophyletic, always forming a grade with *U. neglecta* Jongkind.

**Figure 2.**
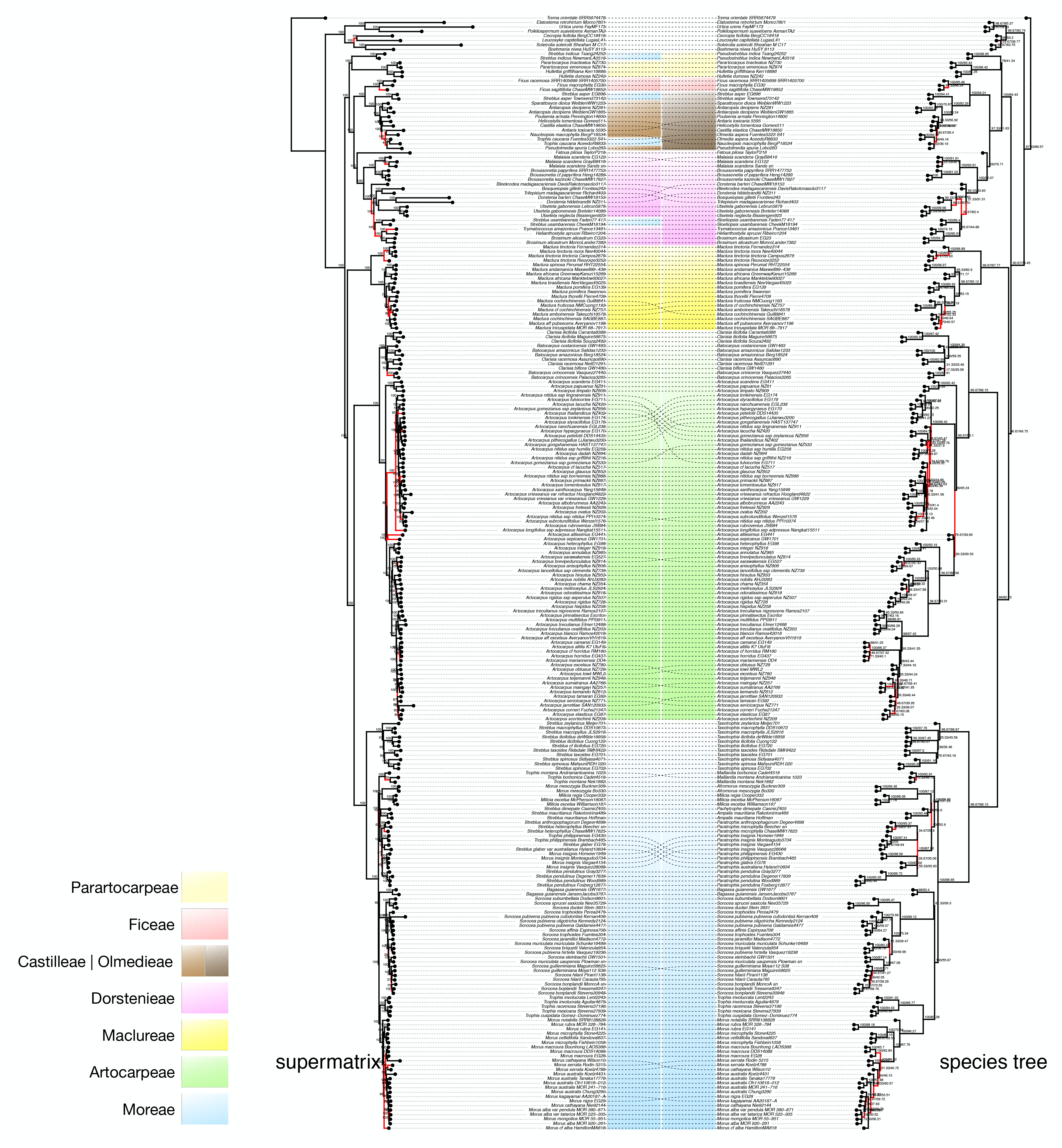
Phylogenetic trees from the “exon” dataset. Maximum-likelihood based on a supermatrix of all loci, with bootstrap support and previous nomenclature (left), and a species tree based on gene trees from all loci with bootstrap/LPP support and revised nomenclature (right). Discordant branches are colored in red, and tribal classifications are shaded without (left) and with (right) the revised classifications presented here.

**Figure 3.**
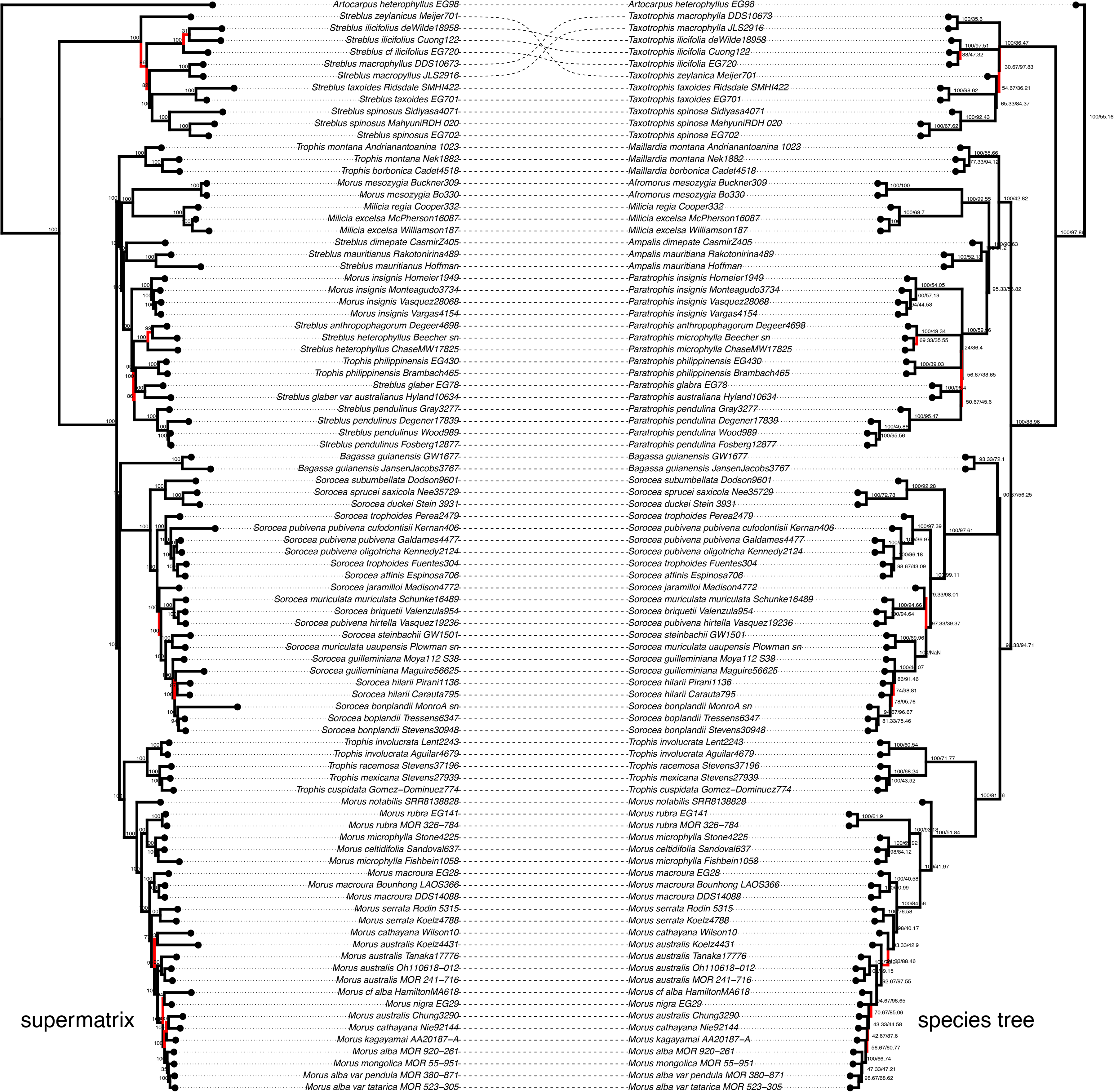
Phylogenetic trees of the Moreae clade from the “supercontig” dataset. Maximum-likelihood based on a supermatrix of all loci, with bootstrap support and previous nomenclature (left), and a species tree based on gene trees from all loci with bootstrap/LPP support and revised nomenclature (right). Discordant branches are colored in red.

The supermatrix and species trees were broadly concordant except within *Artocarpus*, where the species tree was much more consistent with previous analyses within that genus (Gardner et al., in review) (Figs. 2, 3). Otherwise, most differences were at shallow phylogenetic depths, such as relationships within *Streblus* section *Paratrophis*. For both data sets, the polytomy hypothesis was rejected (*P* < 0.05) for Moraceae and all tribe and genus-level clades (following the revised classification presented below) (Figs. 2, 3).

Of the genera in Moreae sensu Clement & Weiblen, only *Milicia* and *Sorocea* were monophyletic; *Morus*, *Streblus*, and *Trophis* were not; *Bagassa* is monotypic and resolved as sister to Sorocea, with which it shares straight stamens. Four Moreae species resolved with other tribes: *Trophis caucana* was nested within Castilleae, *Streblus asper* (Retz.) Lour. was sister to Castilleae, *Streblus indicus* (Bureau) Corner was sister to Parartocarpeae, and *Streblus usambarensis* was nested within Dorstenieae. Moreae was otherwise monophyletic, comprising the following subclades: (1) *Streblus* section *Taxotrophis*, sister to all other Moreae; (2) (a) *Trophis* section *Maillardia*, (b) *Milicia* + *Morus* subgenus *Afromorus*, (c) *Streblus* section *Ampalis*, (d) *Streblus* section *Paratrophis* in part + *Morus* subgenus *Gomphomorus*, (e) *Streblus* section *Paratrophis* in part + *Trophis* section *Calpidochlamys*; (3) *Bagassa* + *Sorocea*; (4) *Trophis* sections *Trophis* and *Echinocarpa* C.C. Berg + *Morus*. These are roughly geographic clades: (1) Southeast Asia; (2) (a) Madagascar, (b) Southeast Africa, (c) Madagascar, (d) Pacific + South America, (e) Southeast Asia + Pacific; (3) South America; (4) South America.

While *Trophis philippinensis* was inside the *Paratrophis* clade in all analyses, the position of *Morus insignis* was not stable. In the “exon” analyses, it was always in *Paratrophis* (Fig. 2), although the polotomy hypothesis could not be rejected for its position as sister to *S. anthropophagorum+S. heterophyllus*. In the “supercontig” and chloroplast analyses, it was sister to *Paratrophis*, and the polotomy test was rejected for that node (Figs. 3, S1). In the five phylogenetic networks reported (Fig. 4, Table S3), *M. insignis* was inside *Paratrophis* in three of them, had a hybrid origin involving *Paratrophis* in one, and was only unambiguously sister to *Paratrophis* in one network, although even in that one it was involved in a hybridization involving *Paratrophis*.

**Figure 4.**
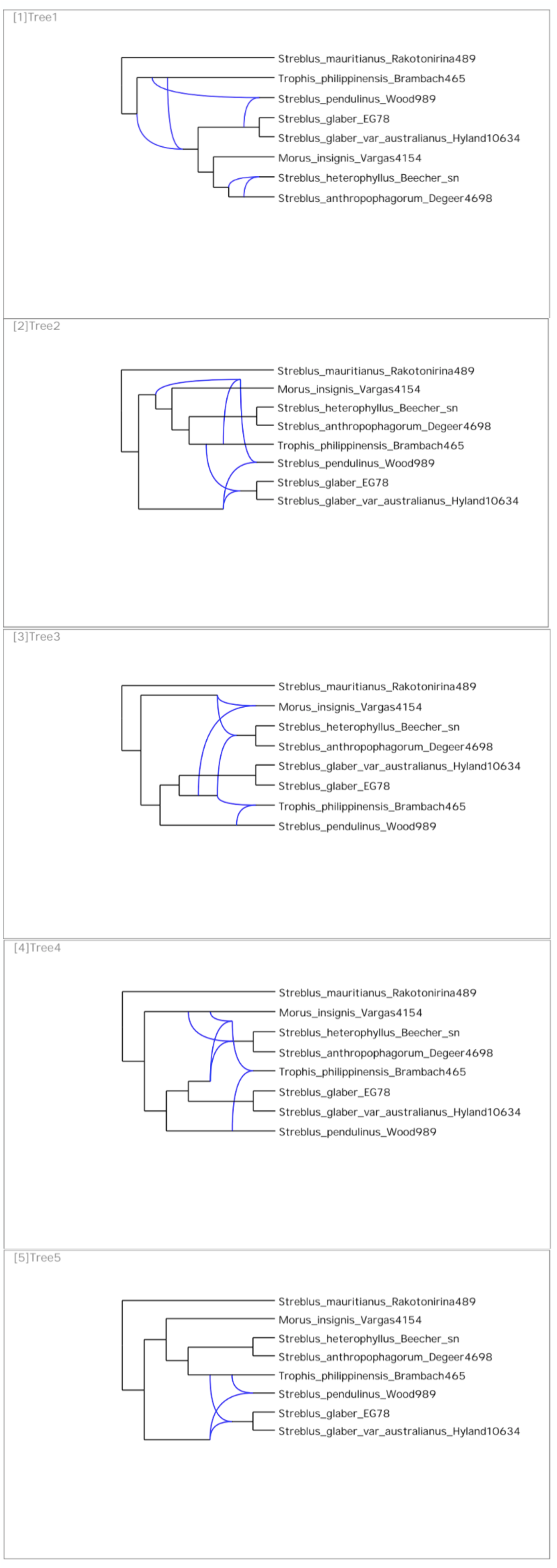
The best five maximum-pseudo-likelihood phylogenetic networks for the *Paratrophis* clade.

The chloroplast phylogenetic tree (Fig. S1) was generally in agreement with the nuclear phylogenetic trees, with three notable differences. *Streblus indicus* and the Parartocarpeae formed a grade paraphyletic to Dorstenieae, Castilleae, and Ficeae, rather than a clade. *Streblus zeylanicus* (Thwaites) Kurz was sister to *S. taxoides* (B. Heyne ex Roth) Kurz (instead of sister to all of section *Taxotrophis*), although with low support (62%), and *S. glaber* subsp. *australianus* was not sister to subsp*. glaber*, although its placement was not strongly supported (80%). Finally, *Morus insignis* was sister to *Paratrophis*, agreeing with the “supercontig” analysis but not the “exon” analysis.

The ITS and *rbc*L phylogenetic trees based on few characters, were not very well resolved, with only few nodes attaining 100% bootstrap support (Fig. S2). Nevertheless, with the exception of a few stray taxa (one *Dorstenia* sample in ITS and one *Streblus indicus* samples in *rbc*L), both phylogenetic trees reconstructed monophyletic tribes with at least moderate support. In the ITS phylogenetic tree, *Streblus smithii* and *S. banksii* were both in a well-supported clade comprising section *Paratrophis*, and *S. tonkinensis* was part of the Castilleae clade, which also contained *S. asper* and *Trophis caucana*. In the *rbc*L phylogenetic tree, the two *Streblus ascendens* samples, the four samples of the undetermined *Streblus* from Papua New Guinea, and *S. smithii* were part of a well-supported clade comprising section *Paratrophis*.

### Time-calibration and ancestral state reconstruction

The cross-validation criterion was minimized by a smoothing parameter (λ) value of 10_6_. The tree calibrated under the correlated model (logLik = −17; p-logLik = −17; ΦIC = 1108) was less sensitive to changes in λ and generally had younger ages than the tree calibrated under the relaxed model (logLik = −56.7; p-logLik = −53129706, ΦIC = 4209013), the latter of which had perhaps implausibly long terminal branches (Table 2, Fig. S3). The crown age of Moreae was Paleocene (59.1 Ma) under the correlated model and late Cretaceous (75.4 Ma) under the relaxed model. Because the correlated tree was more consistent with past family-wide studies (Zerega & al., 2005; Zhang & al., 2019), we used that tree for all further analyses (Fig. 5). The BAMM analysis found a single credible rate shift, not surprisingly at the crown node of *Ficus* (Fig. S4); that rate shift was—also not surprisingly—associated with a loss of inflexed stamens, which are never found in *Ficus* (*P* = 0.035); no rate shifts were found within Moreae. Ancestral reconstruction on the entire tree found that ancestral Moraceae had stamens inflexed in bud; these were lost nine times (in Ficeae, Olmedieae [=Castilleae], Parartocarpeae, twice in Dorstenieae, *Maclura* section *Cudrania* (Trécul) Corner, Artocarpeae, *Bagassa,* and *Sorocea*) and regained once (in *Trophis caucaua*) (Fig. 6). Model testing on both the whole tree and on the Moreae+Artocarpeae+Maclureae subtree indicated that a BiSSE model was not substantially better than a trait-independent model (Moreae only: AIC for BiSSE = 1110.4; trait-independent = 1112.8; ΔAIC = 2.4; whole family: AIC for BiSSE = 1488.3; trait-independent = 1490.4; ΔAIC = 2.1). In any event, the reconstructions were identical (Figs. 6).

**Table 2.**
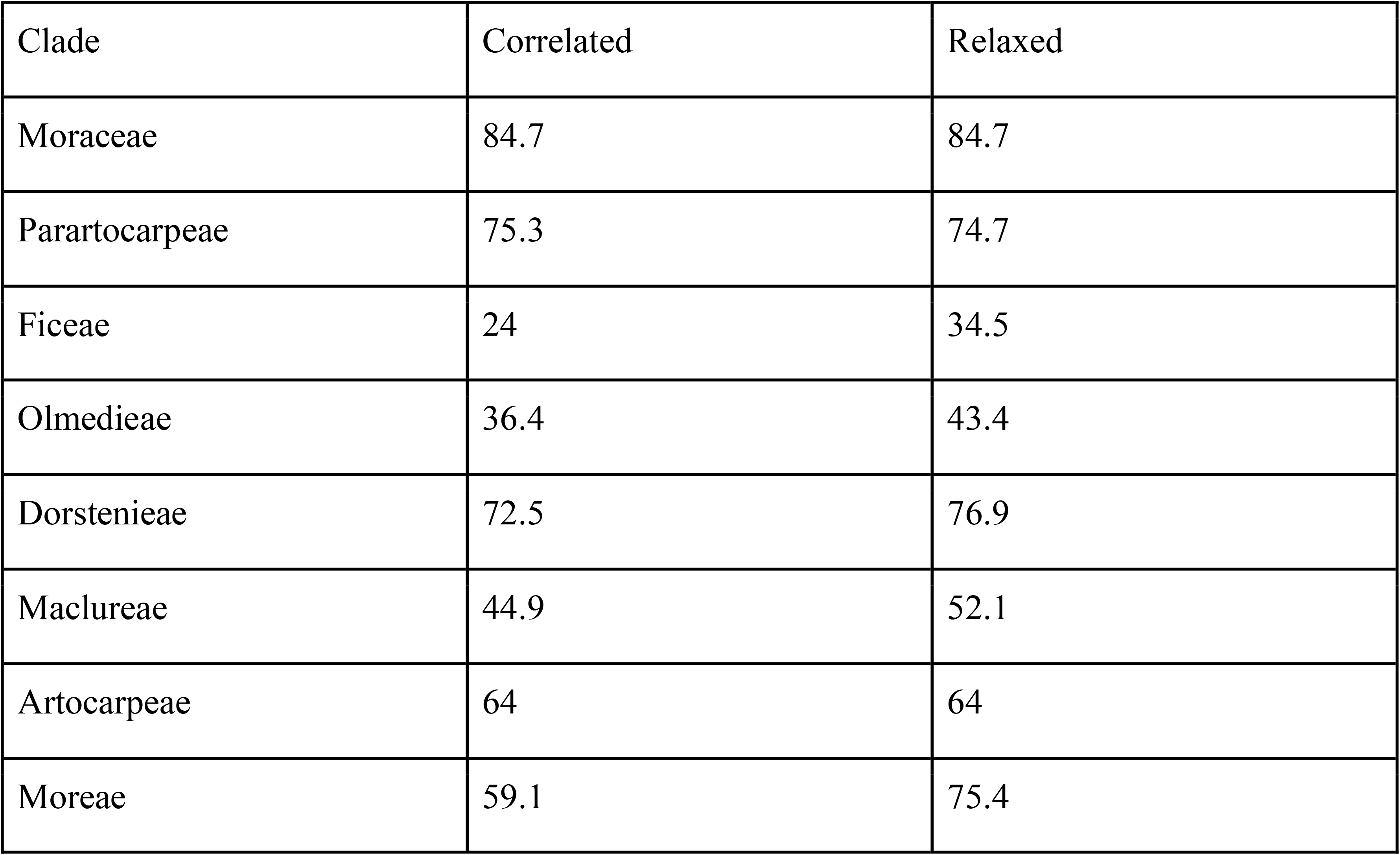
Divergence times (in Ma) estimated using penalized likelihood under correlated and relaxed models. Nomenclature follows the revisions proposed in this study.

| Clade | Correlated | Relaxed |
| --- | --- | --- |
| Moraceae | 84.7 | 84.7 |
| Parartocarpeae | 75.3 | 74.7 |
| Ficeae | 24 | 34.5 |
| Olmedieae | 36.4 | 43.4 |
| Dorstenieae | 72.5 | 76.9 |
| Maclureae | 44.9 | 52.1 |
| Artocarpeae | 64 | 64 |
| Moreae | 59.1 | 75.4 |

**Figure 5.**
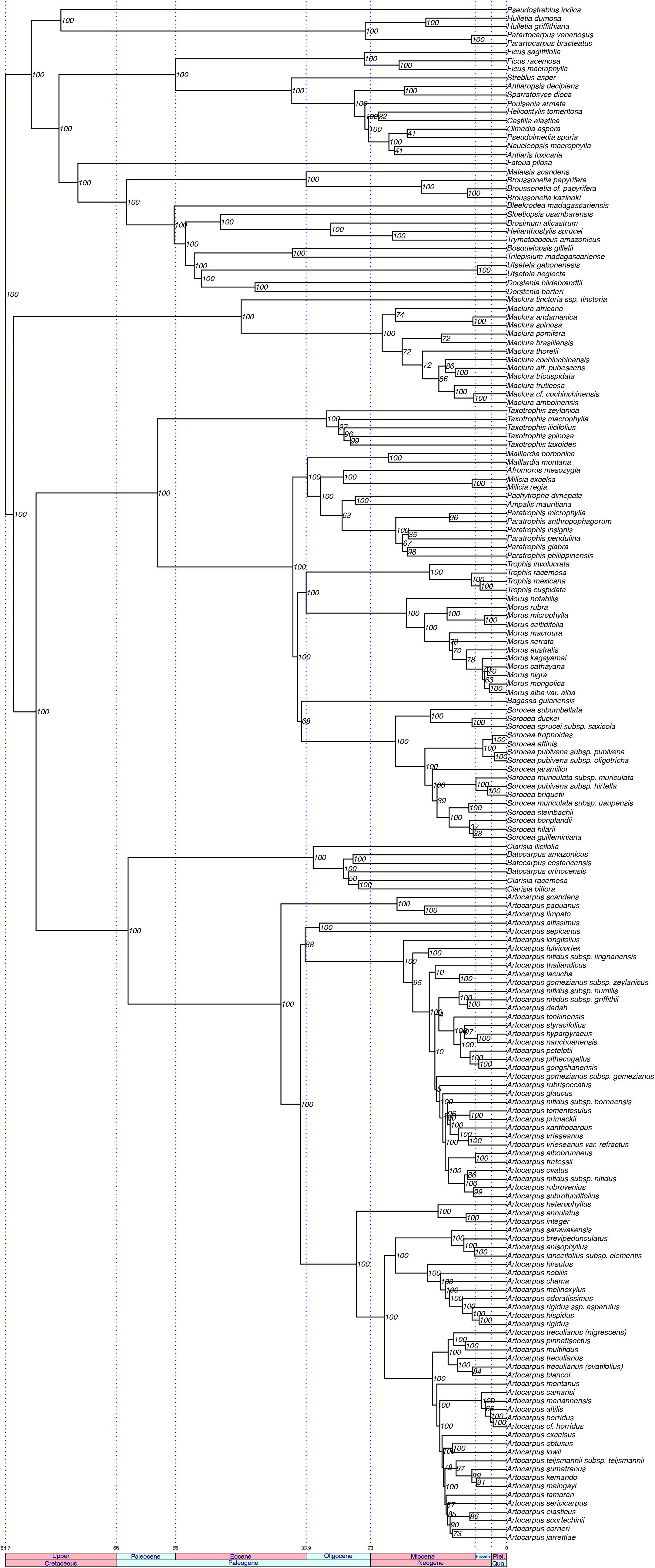
Time-calibrated phylogenetic tree, with revised nomenclature.

**Figure 6.**
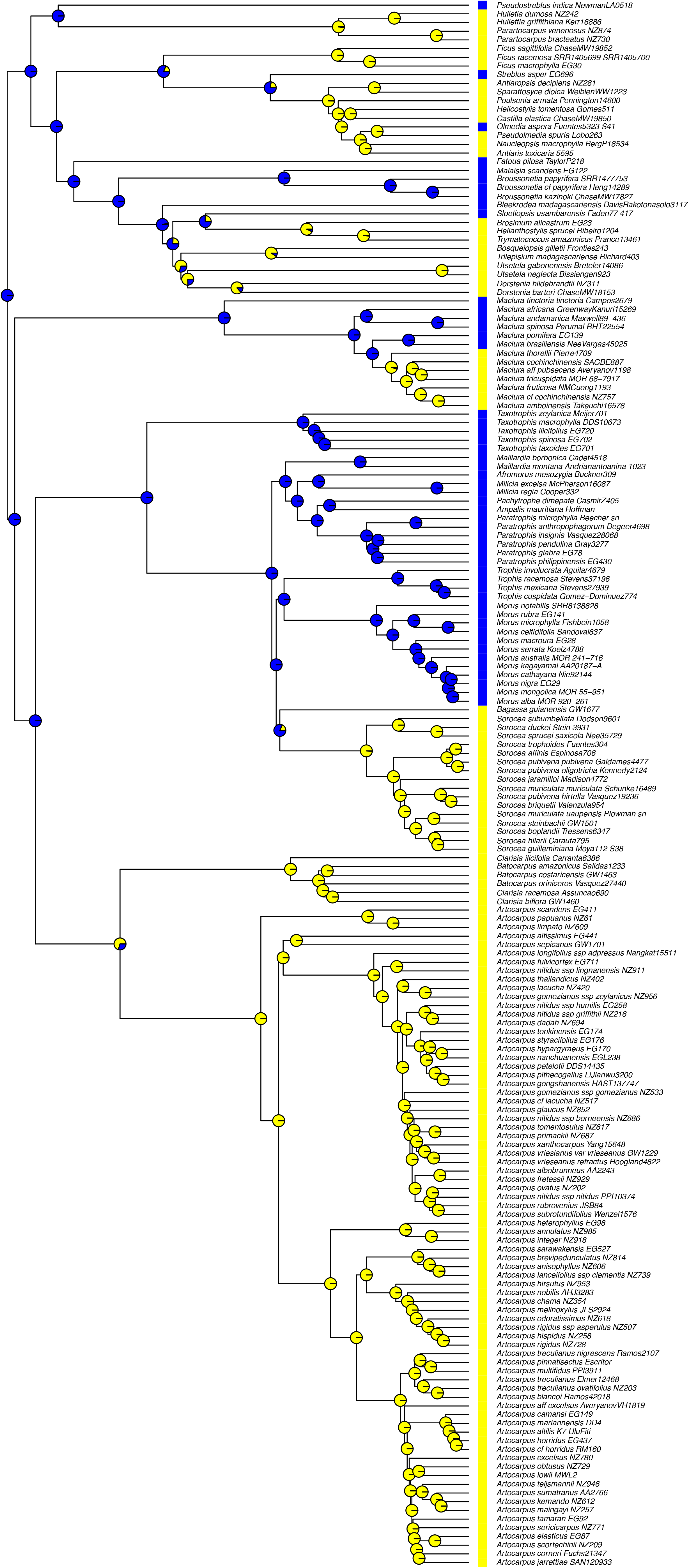
Ancestral reconstruction of stamen position, with revised nomenclature. The reconstruction was identical under a trait-dependent (BiSSE) or a trait-independent model. Blue = inflexed in bud; yellow = straight in bud.

## Discussion

### Sequencing and combination of data sets

This study was materially improved by our ability to combine samples enriched with two largely non-overlapping bait sets. Despite minimal by-design overlap, we were able to assemble many overlapping loci for samples with moderate to deep coverage (Table S2), adding taxa that would not otherwise have been included in the study and replicating taxa to confirm unexpected phylogenetic placement (e.g, *Streblus indicus*). Taxa replicated across the two data sets performed well in phylogenetic analyses, with 15 out of 19 always resolving as monophyletic in supermatrix analyses and 18/19 so resolving in ASTRAL analyses, with only *Utsetela gabonensis* always forming a grade instead (with a difficult-to-distinguish congener). Our results should embolden others to combine and repurpose data sets in similar ways.

### Higher taxonomy in Moraceae and the delimitation of the tribe Moreae

Our results support Corner’s overall approach to the classification of Moraceae, if not all of its details, including the primacy of inflorescence architecture and the unreliability of inflexed stamens for higher taxonomy (Corner, 1962). Inflexed stamens are a plesiomorphy that was lost nine times in Moraceae (Fig. 6), a preserved ancestral character whose past taxonomic importance accounts for the rather extreme non-monophyly of the Moreae. *Streblus* s.l. provides the best illustration of this principle, appearing in four out of seven tribe-level clades within Moraceae. We may justly call its disparate sections, as Corner did, “fragments of an ancestral *Streblus*.” (Corner, 1975)—or in modern parlance, a paraphyletic remnant preserving plesiomorphic staminate flowers similar to those likely to have occurred in the ancestor of all Moraceae, with four free tepals and four inflexed stamens (Clement & Weiblen, 2009). If one of the goals of modern systematics is to establish taxonomic frameworks that reflect as far as possible real evolutionary relationships, our results may serve as a warning to carefully investigate whether characters used for taxonomy are derived (synapomorphic) or ancestral (symplesiomorphic).

Broadly speaking, a tribal delimitation based on inflorescence architecture (supplemented with other characters in some cases) agrees best with the phylogenetic trees presented here. Moreae (as revised below) have unisexual spicate or racemose inflorescences (the pistillate ones sometimes uniflorous), with the globose-capitate pistillate inflorescences of the monotypic *Bagassa* as the sole exception. Stamens may be either inflexed or straight. Elsewhere in the family, racemes and spikes are rare, with the former found in *Maclura* (in part) and the latter found in the related genera *Broussonetia*, *Allaeanthus* Thwaites, and *Malaisia* and arguably in *Batocarpus* H. Karst. and *Clarisia*; all of these have capitate pistillate inflorescences. The taxa of *Streblus* s.l. and *Trophis* s.l. that must be excluded from Moreae all have inflorescences that do not fit our general rule: discoid-capitate (*Streblus asper* and *Trophis caucana*), cymose (*Streblus indicus*), or bisexual (*Streblus usambarensis*). In these cases, perhaps Corner did not take his emphasis on inflorescence architecture quite far enough, including too wide of a variety in this one tribe. In critiquing the utility of inflorescence architecture for classification, Berg (1977b) noted the similarity in the inflorescence structure of *Bleekrodea* (then part of Moreae, and included in *Streblus* by Corner) to that of *Utsetela* and *Helianthostylis* (Dorstenieae), an observation that proved prescient when *Bleekrodea* was found to belong to Dorstenieae (Clement & Weiblen, 2009).

Olmedieae (as revised below, including Castilleae) can be defined entirely based upon the presence of a discoid inflorescence subtended by an involucre of imbricate bracts. Two species with such involucres previously classified as Moreae, *Streblus asper* and *Trophis caucana* (=*Olmedia aspera*) always appeared in the Olmedieae (=Castilleae) clade in our analyses (Figs. 2–3). *Olmedia aspera* (=*Trophis caucana*)—the nomenclatural type of the tribe Olmedieae—was transferred to Moreae because of its inflexed stamens and lack of self-pruning branches (Berg, 1977b). In inflorescence morphology, however, *T. caucana* closely resembles other Castilleae, always subtended by an involucre of imbricate bracts (Berg, 1977b, 2001). The staminate inflorescences of *Streblus asper* are strikingly similar to those of *T. caucana*, and it is remarkable that the affinity between *S. asper* and the Olmedieae has not been seriously considered until now. Previous barcoding or phylogenetic studies have placed *Trophis caucana* (Kress & al., 2009) and *Streblus tonkinensis* (Chen & al., 2016) (closely allied to *S. asper*) in the Castilleae clade, but those results went unremarked upon, perhaps because of the broad scale of the studies (respectively, forest community phylogenetics and the vascular plants of China).

The remaining five tribes can all be broadly defined based on inflorescence architecture as Corner argued, sometimes supplemented by other characters as Berg preferred, allowing of course for the exceptions made inevitable by the vicissitudes of evolution. Ficeae of course is defined by the syconium (essentially an urceolate disc that has been closed at the top). Maclureae have densely-packed globose infructescences and can be distinguished from Artocarpeae by their four stamens (inflexed in most sections) and armature. Artocarpeae also have densely-packed globose infructescences but only one straight stamen and no armature. Parartocarpeae have a few connate involucral bracts and (in large part) flowers embedded in fleshy receptacles. Dorstenieae are perhaps the most heterogenous group, but in large part they have bisexual inflorescences, often capitate or discoid, and often with ballistically-ejected endocarps.

### The role of inflexed stamens in the evolution of Moraceae

The repeated losses of inflexed stamens (Fig. 6), which are associated with wind pollination (Bawa & Crisp, 1980; Berg, 2001), raise the possibility that transitions from wind to animal pollination, which have already been documented in Moraceae (Momose & al., 1998; Sakai & al., 2000; Datwyler & Weiblen, 2004; Gardner & al., 2018), are even more common within the family. Generally considered rare (Culley & al., 2002), the shift from wind to animal pollination may be a repeated feature of Moraceae deserving of further investigation. Further investigation of *Sorocea*, with its straight stamens and sometimes-scented inflorescences, may reveal that, like *Artocarpus*, it contains both wind and animal pollination. And while little is known about pollination in the Dorstenieae, the presence of unisexual inflorescences and inflexed stamens (e.g., *Broussonetia*) as well as bisexual inflorescences with straight stamens (e.g., *Brosimum, Dorstenia*) raises the possibility of transitions within that clade as well; taxa with inflexed stamens but bisexual inflorescences such as *Bleekrodea* and *Sloetia* (which is visited by bees, EMG pers. obs.) may represent remnants of an intermediate state. Finally, the conclusion that a single transition to insect pollination preceded the split between the Ficeae and Olmedieae (Castilleae) should be reevaluated in light of the position of *Streblus asper* as sister to the latter (Figs. 2, 3). The only shift in diversification rates our analyses recovered was on the branch leading to the very diverse *Ficus*, suggesting that the shift away from wind pollination may not by itself lead to increased diversification.

### Explanation of taxonomic revisions

We present a generic revision of Moreae (Fig. 7) based on the present phylogenetic study as well as morphological characters. Arranging monophyletic and morphologically coherent genera requires one change of rank and 8 new combinations, but no entirely new names. These revisions provide a framework for within-genus revisionary work, which will require more intensive sampling and review of specimens within the genera circumscribed here. We provide an explanation of the taxonomic changes, followed by a formal presentation of the affected tribes and genera, with complete species lists and synonymy.

**Figure 7.**
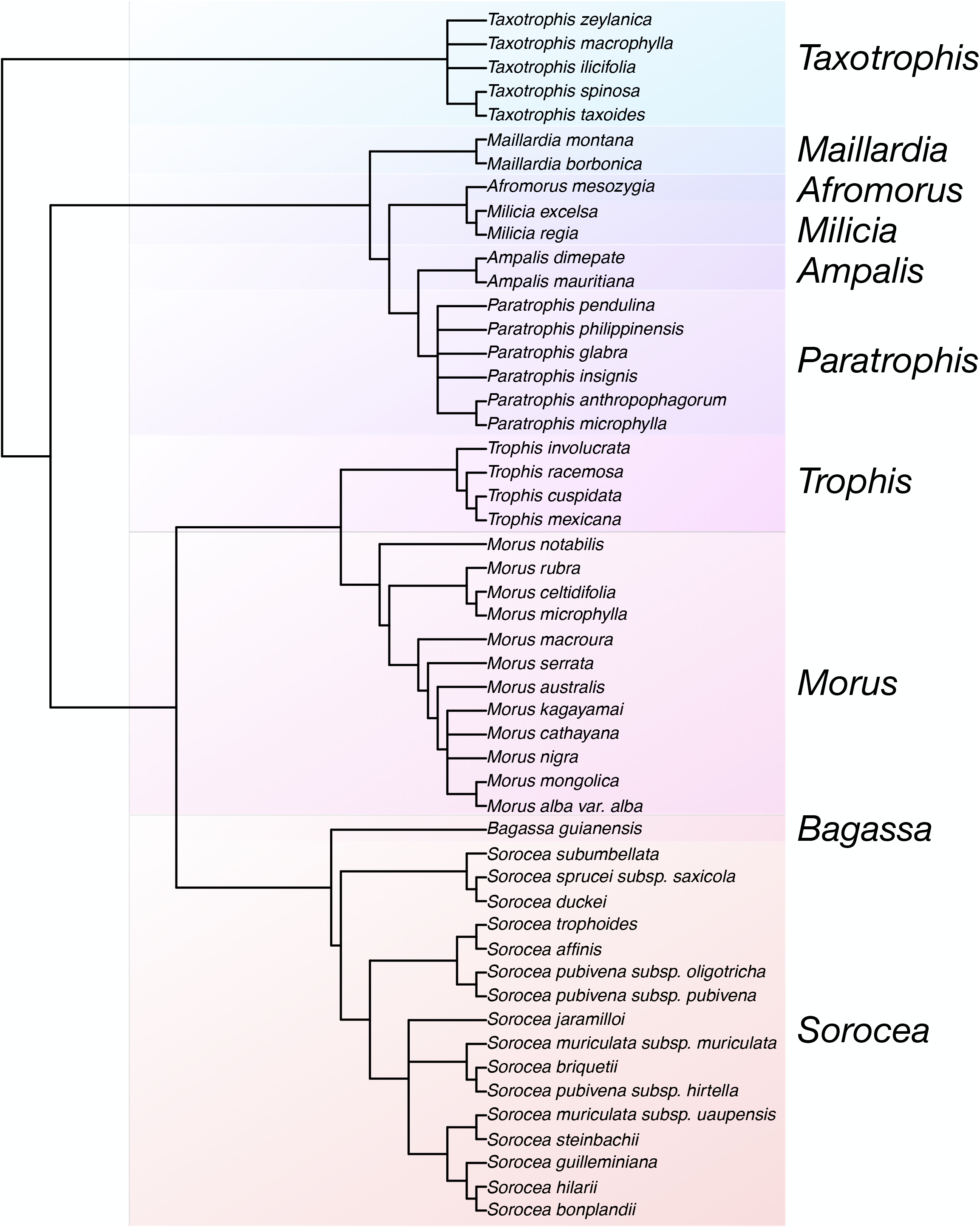
Revised classification of Moreae on a strict consensus tree of all four main phylogenomic analyses.

*Trophis* and the reinstatement of the Olmedieae *—* To make *Trophis* monophyletic, we propose that it be applied strictly to the Neotropical clade that includes the type species *Trophis americana* L. (=*Trophis racemosa* (L.) Urb.). To accomplish this, we reinstate the genus *Maillardia* and transfer *Trophis philippinensis* to *Paratrophis* (discussed below), reinstating its former name *Paratrophis philippinensis* (Bureau) Fern.-Vill. Because *Trophis caucana* does not belong in *Trophis* or Moreae but rather with the members of Castilleae, we reinstate its former name *Olmedia aspera* Ruiz & Pav. and transfer it to that tribe, whose former name Olmedieae must now be reinstated on account of its priority.

*Streblus* — We restrict the genus *Streblus* to three species comprising most of section *Streblus*—*Streblus asper, Streblus tonkinensis*, and *Streblus celebensis* C.C. Berg—and transfer the genus to Olmedieae. Although *S. celebensis* was not included in our phylogeny, the sub-involucrate inflorescences are similar to those of *S. asper*, with which *S. celebensis* differs primarily in vegetative characters, the latter having broadly-toothed margins in the distal half of the leaf. Both occur in Sulawesi, where at least one specimen with intermediate leaf morphology has been collected (Sulawesi, Kendari: *Kjellberg 452*, 24 Feb. 1929, L, det. *S. asper* by E.J.H. Corner). We therefore retain its taxonomic position in *Streblus*. The remaining member of section *Streblus*, *S. usambarensis*, belongs with Dorstenieae, and we therefore transfer it to that tribe, reinstating its former name *Sloetiopsis usambarensis*. *Streblus indicus* is sister to Parartocarpeae, and we therefore reinstate its former name *Pseudostreblus indicus* and transfer it to Parartocarpeae.

The remaining species of *Streblus* s.l. are properly placed in Moreae but are still paraphyletic. We therefore reinstate the genera *Ampalis*, *Paratrophis*, and *Taxotrophis*, largely corresponding to the former sections but requiring some new combinations.

*Paratrophis* as presented here is united by spicate male inflorescences (usually) with peltate or reniform bracts (except for *Paratrophis philippinensis* (=*Trophis philippinensis*), which does not have fleshy tepals in fruit). Within *Paratrophis* we include *Paratrophis ascendens* (Corner) E.M. Gardner (=*Streblus ascendens* Corner) based on its inflorescence morphology and phylogenetic position in the *rbc*L tree (Fig. S2b). Its previous position within the monotypic section *Protostreblus* was due to the type specimen’s spiral phyllotaxy, but the latter may be atypical, as a more recent collection *Womersley NGF 24791* (K, L, BO), has distichous leaves. Berg (1988), recognizing the close affinity between *Pachytrophe dimepate* Bureau and *Ampalis mauritiana*, included both in *Streblus* section *Ampalis*. Our phylogenetic results support this grouping, which we maintain in the reinstated genus *Ampalis*, requiring one new combination. We follow Baillon in maintaining *Pachytrophe* as a section of *Ampalis* in order to recognize the differences between them in phyllotaxy, stipule amplexicaulity, and embryo characters. Further intensive study of that species complex may of course warrant a different approach, including potentially reducing both to a section of *Paratrophis*. Our species lists within the former components of *Streblus* follow Berg’s approach (1988, 2006), with two exceptions. We provisionally recognize *Taxotrophis zeylanica* (Thwaites) Thwaites as distinct from *Taxotrophis taxoides* (B. Heyne ex Roth) W.L. Chew ex E.M. Gardner (=*Streblus taxoides* (B. Heyne ex Roth) Kurz) based on our phylogenetic work and following the *Flora of China*, which recognizes *Streblus zeylanicus* as distinct from *S. taxoides* based on its clustered pistillate inflorescences. In addition, we provisionally recognize *Paratrophis australiana* (=*S. glaber* subsp. *australianus*) as distinct from *Paratrophis glabra* based on its geographic and consistent morphological distinctiveness. *Taxotrophis* and *Paratrophis* warrant further investigation to refine species limits.

*Morus — Morus* has never been revised, and the species concepts are often based on minor morphological differences (Berg et al., 2006; Berg 2001), and the paraphyly of several species in our analyses suggests that a broad *M. alba* similar to Bureau’s (1873) may be worth a second look. The monotypic subgenus *Afromorus* is sister to *Milicia*, but the leaf morphology is markedly different, instead resembling other *Morus* species in its trinerved based and crenate margins. We therefore raise *Afromorus* to genus level, requiring one new combination. *Morus insignis*, from western Central and South America, bears a remarkable resemblance to *Paratrophis*, in particular *P. pendulina*, especially in leaf morphology, which in *M. insignis* is not consistently trinerved as in other mulberries. The infructescence appears superficially like a mulberry because of its basally fleshy tepals, although with more loosely-packed flowers, but closer inspection places it firmly within *Paratrophis*, with drupes protruding from the persistent tepals, peltate bracts, and a sterile groove. Because analyses differ as to whether *M. insignis* is sister to *Paratrophis* or part of it, it seems best to treat them as congeners.

These changes result in ten monophyletic genera of Moreae, providing a framework for revisionary work within the genera. Below, we present a complete genus and species list for Moreae with brief descriptions for all genera and synonymies for new combinations and reinstated or recircumscribed taxa. All cited protologues were reviewed, and dates for works published piecemeal were confirmed by reference to Taxonomic Literature 2 online (https://www.sil.si.edu/DigitalCollections/tl-2/index.cfm).

## Taxonomic Treatment

### Key to the tribes of Moraceae

This key follows the tribal circumscription of Clement & Weiblen (2009), as subsequently modified by Zerega & al. (2010), Chung & al. (2017), Zerega & Gardner (2019), and this study.

1. Inflorescence a syconium (urceolate with the opening entirely closed by ostiolar bracts, flowers enclosed at all stages of development) – Ficeae (*Ficus*)
1. Inflorescence not a syconium (capitate, spicate, discoid, or urceolate, but flowers not entirely enclosed at all developmental stages) – 2
2. Inflorescences (at least staminate) with an involucre of imbricate bracts; often with self-pruning horizontal branches (except *Olmedia*, *Poulsenia, Streblus*) – Olmedieae (*Antiaris, Antiaropsis, Castilla, Helicostylis, Maquira, Mesogyne, Naucleopsis, Olmedia, Perebea, Poulsenia, Pseudolmedia, Sparattosyce*, *Streblus*)
2. Inflorescences not involucrate; plants without self-pruning branches – 3
3. Plants woody; dioecious; pistillate inflorescences globose-capitate; spines axillary or terminating short shoots – Chlorophoreae (*Maclura*)
3. Plants woody, herbaceous, or succulent; dioecious or monoecious; pistillate inflorescences various; spines absent or if present, then pistillate inflorescences not globose capitate – 4
4. Trees or shrubs; monoecious; inflorescences unisexual; staminate flowers with one stamen (rarely two) – Artocarpeae (*Artocarpus, Batocarpus, Clarisia*)
4. Trees, shrubs, lianas, herbaceous, or succulent; monoecious or dioecious; inflorescences unisexual or bisexual; staminate flowers with more than one stamen (or if one stamen then dioecious) – 5
5. Trees or shrubs; monoecious; inflorescences unisexual; stamens straight in bud or staminate flowers 5-parted and inflexed in bud – Parartocarpeae (*Hullettia, Parartocarpus, Pseudostreblus*)
5. Trees, shrubs, lianas, herbaceous, or succulent; monoecious or dioecious; inflorescences bisexual or unisexual; stamens straight or inflexed in bud but staminate flowers never 5-parted – 6
6. Trees, shrubs, lianas, herbaceous, or succulent; inflorescences bisexual (or if unisexual then a climber or herbaceous); endocarp body often ballistically ejected from infructescence – Dorstenieae in part (*Allaeanthus* in part*, Bleekrodea, Bosqueiopsis, Brosimum, Broussonetia* in part, *Dorstenia, Fatoua, Helianthostylis, Malaisia, Scyphosyce, Sloetia, Sloetiopsis, Treculia, Trilepsium, Trymatococcus, Utsetela*)
6. Trees or shrubs; inflorescences unisexual; endocarp body never ballistically ejected – 7
7. Trees; stamens inflexed in bud; pistillate inflorescences globose-capitate – Dorstenieae in part (*Allaeanthus* in part*, Broussonetia* in part)
7. Trees or shrubs; stamens inflexed or straight in bud; pistillate inflorescences various, not globose-capitate if stamens are inflexed in bud – Moreae (*Afromorus, Ampalis, Bagassa, Maillardia, Milicia, Morus, Paratrophis, Taxotrophis, Sorocea, Trophis*)

### Revisions

#### Tribe MOREAE

**Moreae** Gaudich. in Freyc., Voy. Uranie, Bot. (1830) 509.

Moreae subtribe Soroceae Miq in Martius, Fl. Bras. 4, 1, fasc. 12: 111 (1853), p.t. –

Soroceae (Miq.) C.C. Berg, Blumea 50: 537 (2005), p.p.

Strebleae Bureau in DC., Prodr. 17: 215 (1873), p.p.

Tree or shrubs, monoecious or dioecious. ***Leaves*** alternate or opposite, distichous or spirally arranged; stipules lateral to amplexicaul. ***Inflorescences*** unisexual, uniflorous or racemose, spicate, capitate, or globose; bracteate; tepals 4, free to connate; staminate flowers with 4 stamens, filaments straight or inflexed in bud, pistillode usually present; pistillate flowers with (mostly) free ovary, 2 stigmas. ***Fruits*** drupaceous or achene-like with a fleshy persistent perianth, dehiscent or not. ***Seeds*** with or without endosperm, testa usually with a thick vascularized part below the hilum, cotyledons equal or unequal, straight or folded.

Genera and distribution: ten genera and 63 species with a worldwide distribution.

#### Afromorus

***Afromorus*** E.M. Gardner, gen. nov. — based on *Morus* L. *subg. Afromorus* [A. Chev., Rev. Bot. Appl. Agr. Trop. 29 (315–316): 70 (1949), invalidly published]; J.-F. Leroy, Rev. Bot. Appl. Agr. Trop. 29 (323–324): 482 (1949) & Bull. Mus. Hist. Nat. Paris, ser. 2, 21: 732 (1949). TYPE: *Afromorus mesozygia* (Stapf ex A. Chev.) E.M. Gardner.

Dioecious trees, shoot apices deciduous. ***Leaves*** distichous, triplinerved or at least trinerved at the base. ***Stipules*** free, more or less lateral. ***Inflorescences*** solitary or paired, bracts of varying shapes. Staminate inflorescences spicate, to 2.5mm long, flower 4-parted, tepals imbricate, ciliolate, pistillode small, apiculate. Pistillate inflorescences subglobose, ca. 5 mm across, flowers 4-parted, tepals ciliolate, stigma bifid, equal or unequal, arms filiform to 5 mm long. ***Infructescences*** subglobose or less often slightly elongate, ca. 1 mm across, tepals fleshy, yellowish to greenish, drupes ca. 5 × 3–5 mm. ***Seeds*** ca. 4.5 × 2.5–4.5 mm.

Species and distribution: one species, in tropical Africa.

Note: The leaves of *Afromorus*—with crenate margins and trinerved bases—bear a striking resemblance to those of *Morus*, and it is thus not surprising that the former was heretofore included within the latter. However, leaves of *Afromorus* are distinct in being usually completely triplinerved, without an upper pinnately-veined portion as in most species of *Morus*. As the genus is monotypic, the type must be its only species, *Afromorus mesozygia* (Stapf ex A. Chev.) E.M. Gardner.

The genus name is based on Leroy’s invalidly published *Morus* subgenus *Afromorus*, which was published only in French without a Latin diagnosis or description as required under the Code.

1. ***Afromorus mesozygia*** (Stapf ex A. Chev.) E.M. Gardner, comb. nov. — based on *Morus mesozygia* Stapf ex A. Chev. [Végétaux utiles de l’Afrique tropicale française 5: 263 (1909), nomen solum], J. Bot. (Morot) ser. 2, t. 2: 99 (1909).

*Celtis lactea* Sim, For. Fl. Port. E. Afr. : 97, t. 96 (1909), probably post-dating *Morus mesozygia* (fide Berg. 1977) – *Morus lactea* (Sim) Mildbr., Notizbl. Bot. Gart. Berlin 8: 243 (1922) – *Morus mesozygia* var. *lactea* (Sim) A. Chev., Rev. Bot. Appl. Agr. Trop. 29: 72 (1949).
*Morus mesozygia* var. *sanda* A. Chev., Rev. Bot. Appl. Agr. Trop. 29: 71 (1949), invalidly published (under Art. 39.1, Latin description or diagnosis lacking).
*Morus mesozygia* var. *colossea* A. Chev., Rev. Bot. Appl. Agr. Trop. 29: 71 (1949), invalidly published (under Art. 39.1, Latin description or diagnosis lacking).

#### Ampalis

*Ampalis* Bojer, Hort. Maurit. 291 (1837)

*Streblus* Lour. subgen. *Parastreblus* Blume, Mus. Bot. Ludg.-Bat. 2: 89 (1856)
*Streblus* Lous. sect. *Ampalis* (Bojer) C.C. Berg, Proc. Kon. Ned. Akad. Wetensch. C 91(4): 358 (1988).
*Pachytrophe* Bureau, Prodr. [A.P. de Candolle] 17: 234 (1873). – *Ampalis* Boj. sect.

*Pachytrophe* (Bur.) Baillon, Hist. Pl. 6: 191 (1875–76)

Dioecious trees or shrubs. ***Leaves*** distichous to spirally arranged, pinnately veined. ***Stipules*** free, nearly lateral. ***Inflorescences*** solitary or paired in the leaf axils, spicate, with an abaxial sterile groove, flowers in longitudinal rows, bracts basalt attached to subpeltate. Staminate inflorescences to 9 cm long, flowers 4-parted, decussate-imbricate, stamens 4, inflexed in bud, pistillode present. Pistillate inflorescences to 12 cm long, tepals 4, separate, decussate-imbricate, ovary free, stigmas 2, equal. ***Infructescences*** with enlarged fleshy perianths ca. 6–8 mm long, surrounding drupaceous fruits, the latter ca. 5–6 mm long. ***Seeds*** ca. 4 × 4 mm, testa thickened and not distinctly vascularized, cotyledons equal.

Species and distribution: Two species, native to Madagascar and Comoros.

Note: The two sections differ in their phyllotaxy (usually spiral in *Ampalis* and ditichous in *Pachytrophe*) and stipules (free in *Ampalis*, connate in *Pachytrophe*). Sect. *Ampalis* usually has somewhat larger inflorescences.

*Ampalis* sect. *Ampalis*

1. ***Ampalis mauritiana*** (Jacq.) Urb., Symb. Antill. 8: 165 (1920) – *Morus mauritiana* Jacq.,
  Collect. 3: 206 (“1789”, 1791) *– Streblus mauritianus* (Jacq.) Blume, Mus. Bot.
  Lugd.-Bat. 2: 80 (1856).
  *Streblus maritimus* Palaky, Catal. Pl. Madag. 2: 31 (1907)
  *Morus nitida* Willem. In Useri, Ann. Bot. 18: 56 (1796)
  *M. ampalis* Poir. In Lam., Encycl. Bot. 4: 380 (1797)
  *Trophis cylindrica* Roxb., Fl. Indica 3: 599 (1832), pro syn. *M. mauritianae*.
  *Ampalis madagascariensis*, Bojer, Hort. Maurit. 291 (1837) nom. illeg., pro syn.
  *Ampalis madagascariensis* var. *occidentalis* Léandri, Mém. Inst. Sci. Madag., ser. B, 1: 12, pl. (1948).
  *Morus rigida* Hassk., Cat. Hort. Bog. 74 (1844).
  ***Ampalis* Boj. sect. *Pachytrophe*** (Bureau) Baillon, Hist. Pl. 6: 191 (1875–76) ≡ *Pachytrophe* Bureau in A.DC, Prodr. 17: 234 (1873). – Type: *Pachytrophe dimepate* Bureau.
2. ***Ampalis dimepate*** (Bureau) E.M.Gardner, comb. nov., based on *Pachytrophe dimepate* Bureau in A.DC Prodr. 17: 234 (1873) – *Streblus dimepate* (Bureau) C.C.Berg, Proc. Kon. Ned. Akad. Wetensch. C 91(4): 358 (1988).

*Pachytrophe obovata* Bureau, Prodr. [A.P. de Candolle] 17: 235 (1873).
*Plecospermum bureaui* A.G. Richt., Term. Füzetek 18: 296 (1895), nomen nudum et superfl., pro. syn. nom. ined. “Plecospermum obovatum Bureau” (*Boivin 1717*, P).
*Plecospermum (?) laurifolium* Baill. in Grandidier, Hist. Madag. Vol. 35, Hist. Nat. Pl., Tome 5, Atlas 3, 2e partie: pl. 294a (1895) – *Pachytrophe obovata* var. *laurifolia* (Baill.) Léandri, Mém. Inst. Sci. Madag., ser. B, 1: 16 (1948).
*Pachytrophe obovata* var. *montana* Léandri, Mém. Inst. Sci. Madag., ser. B, 1: 16 (1948).

#### Bagassa

*Bagassa* Aubl., Pl. Guiane. Suppl. 15 (1775).

Dioecious trees. ***Leaves*** opposite and decussate, lamina triplinerved, 3-lobed to entire. ***Stipules*** free, lateral. ***Inflorescences*** solitary or paired in the leaf axils, bracteate. Staminate inflorescences spicate with an abaxial sterile groove, to 12 cm long, flowers in longitudinal rows with, tepals 4, stamens 2, straight in bud, pistillode present. Pistillate inflorescences globose-capitate, ca. 1–1.5 cm across, flowers 4-lobed to 4-parted, stigmas 2, filiform. ***Infructescences*** globose, ca. 2.5–3.5 cm across, green, tepals fleshy, yellowish to greenish, drupes ca. 7–8 mm long. ***Seeds*** ca. 3 × 2 mm, testa thin and not vascularized, cotyledons equal.

Species and distribution: One, in tropical South America.

1. ***Bagassa guianensis*** Aubl., Pl. Guiane. Supple. 15 (1775)

#### Maillardia

*Maillardia* Frapp. ex Duch. in Maillard, Notes sur l’île de la Réunion, Annex P. 3 (1862) *Trophis* P. Br. sect. *Maillardia* (Duch.) Corner, Gard. Bull. Singapore 19: 230 (1962).

Dioecious trees or shrubs. ***Leaves*** distichous, pinnately veined. ***Stipules*** free, semi-amplexicaul. ***Inflorescences*** solitary or paired in the leaf axils, bracts subpeltate. Staminate inflorescences spicate with an abaxial sterile groove, flowers 4-parted, decussate-imbricate, stamens 4, inflexed in bud, pistillode present. Pistillate inflorescences up to three together, uni-or bi-florous, perianth tubular 4-lobed, ovary adnate to the perianth, stigmas 2, equal. ***Infructescences*** with enlarged fleshy perianths surrounding drupaceous fruits, the latter to 18 mm long. ***Seeds*** to 13 mm long, testa thin, with a thickened vascularized part below the hilum, cotyledons unequal.

Species and distribution: Two species, in Madagascar, Comoros, and the Seychelles.

1. ***Maillardia borbonica*** Duch., Ann. Notes Réunion, Bot. 1: 148 (1862) – *Trophis borbonica* (Duch.) C.C.Berg, Proc. Kon. Ned. Akad. Wetensch. C 91(4): 355 (1988)

*Maillardia lancifolia* Frapp. ex Duch., Ann. Notes Réunion, Bot. 1: 148 (1862).
*Trophis borbonica* (Duch.) C.C. Berg, Proc. Kon. Ned. Akad. Wetensch. C 91: 355 (1988).
2. ***Maillardia montana*** Leandri, Mém. Inst. Sci. Madagascar, Sér. B, Biol. Vég. 1: 25

(1948) – *Trophis montana* (Leandri) C.C.Berg, Proc. Kon. Ned. Akad. Wetensch. C 91(4): 355 (1988).
*Maillardia occidentalis* Leandri, Mém. Inst. Sci. Madagascar, Sér. B, Biol. Vég. 1: 26 (1948)
*Maillardia orientalis* Léandri, Mém. Inst. Sci. Madagascar, Sér. B, Biol. Vég. 1: 27 (1948)
*Maillardia mandrarensis* Léandri, Mém. Inst. Sci. Madagascar, Sér. B, Biol. Vég. 1: 28 (1948).
*Maillardia pendula* Fosberg, Kew Bull. 29: 266, t. 2 (1974).

#### Milicia

*Milicia* Sim, For. Fl. Port. E. Afr. 97 (1909).

Dioecious trees. ***Leaves*** distichous, lamina pinnately veined. ***Stipules*** free, not fully amplexicaul. ***Inflorescences*** spicate, usually solitary in the leaf axils or on leafless nodes, spicate, flowers in longitudinal rows alternating with rows of bracts, bracts mostly basally attached, abaxial sterile groove present. Staminate inflorescences to 20 cm long, flowers 4-parted, imbricate, stamens 4, inflexed in bud, pistillode present. Pistillate inflorescences to 4.5 cm long, flowers 4-parted, decussate-imbricate, ovary free, stigmas 2, unequal. ***Infructescences*** with enlarged fleshy perianths, surrounding slightly flattened drupaceous fruits, the latter to ca. 3 mm long. ***Seeds*** ca. 2 mm long, testa thin with a slightly thickened vascularized part below the hilum, cotyledons equal.

Species and distribution: Two species in tropical Africa.

1. ***Milicia excelsa*** (Welw.) C.C. Berg, Bull. Jard. Bot. Natl. Belg. 52(1-2): 227 (1982) – *Morus excelsa* Welw., Trans. Linn. Soc. Lond. (Bot.) 27: 69, t. 23 (1869) – *Maclura excelsa* (Welw.) Bur. In DC., Prodr. 17: 231 (1873) – *Chlorophora excelsa* (Welw.)

Benth. & Hook., Gen. Pl. 3(1): 363 (1880).
*Chlorophora tenuifolia* Engl., Bot. Jahrb. 20: 139 (1894).
*Milicia africana* Sim, For. Fl. Port. E. Afr. 97, t. 122 (1909).
*Chlorophora alba* A. Chev., Bull. Soc. Bot. Fr. 58 (Mém. 8d): 209 (1912).
2. ***Milicia regia*** (A. Chev.) C.C. Berg, Bull. Jard. Bot. Natl. Belg. 52(1-2): 227 (1982) –

*Chlorophora regia* A. Chev., Bull. Soc. Bot. Fr. 58 (Mém. 8d): 209 (1912) – *Maclura regia* (A. Chev.) Corner, Gard. Bull. Singapore 19: 237 (1962).

#### Morus

*Morus* L., Sp. Pl. 986 (1753).

Dioecious trees or shrubs. ***Leaves*** trinerved (to five-nerved) at the base. ***Stipules*** free, nearly lateral. ***Inflorescences*** solitary or paired in the leaf axils, without an obvious sterile groove, interfloral bracts absent. Staminate inflorescences spicate or racemose, to 8 cm long, flowers 4-parted, imbricate, stamens 4, inflexed in bud, pistillode present. Pistillate inflorescences subcapitate to spicate, up to 16 cm long, flowers tepals 4, separate, decussate-imbricate, ovary free, stigmas 2, equal. ***Infructescences*** with enlarged fleshy perianths enclosing achene-like fruits, the latter ca. 1 mm long. ***Seeds*** less than 1 mm long, cotyledons equal.

Species and distribution: Sixteen species, Asia and North to Central America; introduced worldwide.

1. ***Morus alba*** L., Sp. Pl. 2: 986 (1753).

**var. *alba***
**var. *multicaulis*** (Perrottet) Loudon, Arbor. Frutic. Brit. 3: 1348 (1838).
2. ***Morus australis*** Poir. in Desrousseaux et al., Encycl. 4: 380 (1797).
3. ***Morus boninensis*** Koidz., Bot. Mag. (Tokyo) 31: 38 (1917)
4. ***Morus cathayana*** Hemsl., J. Linn. Soc., Bot. 26: 456 (1894)

**var. *cathayana***
**var. gongshanensis** (Z. Y. Cao) Z. Y. Cao, Acta Bot. Yunnan. 17: 154 (1995).
5. ***Morus celtidifolia*** Kunth, Nov. Gen. Sp. [H.B.K.] 2: 33 (1817).
6. ***Morus liboensis*** S. S. Chang, Acta Phytotax. Sin. 22: 66 (1984).
7. ***Morus macroura*** Miq., Pl. Jungh. 1: 42 (1851).

**var. *macroura***
**var. *laxiflora*** G.K.Upadhyay & A.A.Ansari, Rheedea 20: 44 (2010).
8. ***Morus koordersiana*** J.-F.Leroy, Bull. Mus. Natl. Hist. Nat. sér. 2, 21: 729 (1949) (endemic to Sumatra and possibly synonymous with *M. macroura*. However, it was not cited by Berg et al. (2006) either as a good species or as a synonym of the latter).
9. ***Morus microphylla*** Buckley, Acad. Nat. Sci. Philadelphia 1862: 8 (1863) (recognized in the *Flora of North America* (Flora of North America Editorial Committee, 1993) but likely conspecific with *M. celtidifolia* and considered so by Berg (2001)).
10. ***Morus mongolica*** (Bureau) C. K. Schneid. in Sargent, Pl. Wilson. 3: 296 (1916).
11. ***Morus nigra*** L. Sp. Pl. 2: 986 (1753).
12. ***Morus notabilis*** C. K. Schneid. in Sargent, Pl. Wilson. 3: 293 (1916).
13. ***Morus rubra*** L., Sp. Pl. 986 (1753).

**var. *rubra***
**var. *murrayana*** (Saar & Galla) Saar, Phytologia 91: 106 (2009).
14. ***Morus serrata*** Roxb., Fl. Ind., ed. 1832, 3: 596 (1832).
15. ***Morus trilobata*** (S. S. Chang) Z. Y. Cao, Acta Phytotax. Sin. 29: 265 (1991).
16. ***Morus wittiorum*** Hand.-Mazz., Anz. Akad. Wiss. Wien, Math.-Naturwiss. Kl. 58: 88 (1921)

#### Paratrophis

*Paratrophis* Blume, Ann. Mus. Bot. Lugduno-Batavi 2 (1856) 81.

*Pseudomorus* Bureau, Ann. Sci. Nat., Bot. sér. 5, 11: 371 (1869).

*Uromorus* Bureau in A.DC., Prodr. 17: 236 (1873).

*Calpidochlamys* Diels, Bot. Jahrb. Syst. 67: 172 (1935); — *Trophis* sect.

*Calpidochlamys* Corner, Gard. Bull. Singapore 19: 230 (1962).

*Chevalierodendron* J.-F. Leroy, Compt. Rend. Hebd. Séances Acad. Sci. 227: 146 (1948).

*Streblus* Lour. sect. *Paratrophis* (Blume) Corner, Gard. Bull. Singapore 19: 216 (1962).

*Streblus* Lour. sect. *Protostreblus* Corner, Blumea 18: 393 (1970).

*Morus* L. *subg. Gomphomorus* J.-F. Leroy, Bull. Mus. Hist. Nat. (Paris), Sér. 2, 21: 732 (1949)

Dioecious trees or shrubs. ***Leaves*** distichous (or spiral in *P. ascendens*), pinnately veined, sometimes trinerved at the base. Stipules free, lateral. ***Inflorescences*** axillary, solitary or up to 5 together, spicate, with an abaxial sterile groove, interfloral bracts mostly peltate, flowers sessile in longitudinal rows, tepals 4, valvate, ciliolate. Staminate inflorescences up to at least 20 cm. long, flowers with filaments inflexed in bud, pistillode present. Pistillate inflorescences up to at least 10 cm long, flowers usually at least 2 (to many), tepals free (except in *P. philippinensis*), stigma bifid, arms equal. ***Fruits*** drupaceous, red to black, with tepals persistent but usually not enlarged or fleshy (except in *P. philippinensis* and *P. insignis*), up to ca. 1 cm long. ***Seeds*** up to ca. 8 × 6 mm, cotyledons equal.

Species and distribution: Twelve species from the Malesian region to Australia, New Zealand, and Oceania, Central and western South America.

1. ***Paratrophis anthropophagorum*** (Seem.) Benth. & Hook. f. ex Drake, Ill. Fl. Ins. Pacif. 296 1892

*Trophis anthropophagorum* Seem. [Bonplandia 9 (17–18): 259 (1861), nomen nudum] Fl. Vit. 258, t. 68 (1868); — *Uromorus anthropophagorum* (Seem.) Bureau in de Candolle, Prodr. 17 (1873) 236; — *Streblus anthropophagorum* (Seem.) Corner, Gard. Bull. Sing.19 (1962) 221.
*Caturus oblongatus* Seem., Fl. Vit. 254 (1868).
*Paratrophis ostermeyri* Rech., Fedd. Rep. 5: 130 (1908).
*Paratrophis viridissima* Rech., Fedd. Rep. 5: 130 (1908).
*Paratrophis zahlbruckneri* Rech., Fedd. Rep. 5: 130 (1908).
*Pseudomorus brunoniana* (Endl.) Bureau var. *tahitensis* J. Nadeaud, Énum. Pl. Tahiti 43 (1873) – *Uromorus tahitensis* (J. Nadeaud) Bureau in de Candolle, Prodr. 17: 237 (1873) – *Paratrophis tahitensis* (Bureau) Benth. & Hook.f. ex Drake, Ill. Ins. Mar. Pacif. Fasc. 7: 296 (1892); et Fl. Polyn. Franc. 193 (1892). – *Streblus tahitensis* (J. Nadeaud) E.J.H. Corner, Gard. Bull. Singapore 19: 225 (1962).
2. ***Paratrophis ascendens* (Corner) E.M. Gardner comb. nov.** — based on *Streblus ascendens* Corner, Blumea 18: 395, t.1 (1970).
3. ***Paratrophis australiana*** C.T. White, Contr. Arnold Arbor. 4: 15 (1933) — *Streblus glaber* (Merr.) Corner var. *australianus* (C.T. White) Corner, Gard. Bull. Singapore 19: 221 (1962) — *Streblus glaber* (Merr.) Corner subsp. *australianus* (C.T. White) C.C. Berg, Blumea 50: 548 (2005).
4. ***Paratrophis banksii*** Cheeseman, Man. N.Z. Fl. 633 (1906) – *Streblus banksii* (Cheeseman) C.J. Webb, in Connor & Edgar, New Zealand J. of Bot. 25 136 (1987).

*Paratrophis heterophylla* var. *elliptica* Kirk T.N.Z.L 29: 500, t. 46 (1897); — *Streblus heterophyllus* var. *ellipticus* (Kirk) Corner, Gard. Bull. Singapore 19: 222 (1962). Note:— Morphologically very close to and not always distinguishable from *P. microphylla*.
5. ***Paratrophis glabra*** (Merr.) Steenis, J. Bot. 72: 8 (1934) – *Gironniera glabra* Merr., Philipp. J. Sci., 1, Suppl. 42 (1906); Enum. Philipp. Flow. Pl. 2: 35 (1923) — *Chevalierodendron glabrum* (Merr.) J.-F. Leroy, Compt. Rend. Hebd. Séances Acad. Sci. 227: 146 (1948) — *Streblus glaber* (Merr.) Corner, Gard. Bull. Singapore 19: 221 (1962). *Aphananthe negrosensis* Elmer, Leafl. Philipp. Bot. 2: 575 (1909). *Pseudostreblus caudatus* Ridl., J. Fed. Malay States Mus. 6: 54 (1915). *Streblus laevifolius* Diels, Bot. Jahrb. Syst. 67: 171 (1935). *Streblus urophyllus* Diels, Bot. Jahrb. Syst. 67: 172 (1935) — *Streblus glaber* (Merr.) Corner subsp. *urophyllus* (Diels) C.C. Berg, Blumea 50: 548 (2005); *Streblus urophyllus* Diels var. *salicifolius* Corner, Gard. Bull. Singapore. 19: 225 (1962). Note: Berg et al. (2006) treated *P. australiana* and *Streblus urophyllus* as subspecies of *Streblus glaber*. *Paratrophis australiana*, endemic to Australia, has crenate leaf margins and somewhat larger staminate inflorescences with more interfloral bracts than *P. glabra*; these morphological differences are consistent and geographically confined, and we therefore reinstate *P. australiana*. This stands in contrast to *Streblus urophyllus*, which we provisionally maintain in synonomy under *P. glabra*. Although collections from Mt. Wilhelm in New Guinea are remarkable for their thick coriaceous leaves and spinose margins, similar toothed margins can be found at higher elevations in Borneo and Sulawesi, suggesting that the striking leaf morphology of *S. urophyllus* is an alpine effect not indicative of speciation. We reserve judgment on the status of *Streblus urophyllus* var. *salicifolius* Corner, applied by Corner (1962) to specimens with linear leaves, provisionally maintaining it in synonomy following Berg et al. (2006).
6. ***Paratrophis insignis* (Bureau) E.M. Gardner comb. nov.** — based on *Morus insignis*

Bureau, in De Candolle, Prodr. 17: 247 (1873).
*Morus peruviana* Planch. ex Koidz., Fl. Symb. Orient.-Asiat. 88 (1930).
*Morus trianae* J.-F.Leroy, Bull. Mus. Hist. Nat. (Paris) Sér. 2, 21: 731 (1949).
*Morus marmolii* Legname, Lilloa 33: 334 (1973) Note: The infructescences differ from most other *Paratrophis* in their fleshy free tepals and denser aggregation of flowers, but the staminate inflorescences are typical of the genus.
7. ***Paratrophis microphylla*** (Raoul) Cockayne, Bot. Notes Kennedy’s Bush & Scenic Res.

Port Hills, Lyttelton (Rep. Scenery Preserv.) 3 (1915) – *Epicarpurus microphyllus*
Raoul, Choix Pl. Nouv.-Zel. 14. (1846) – *Taxotrophis microphylla* (Raoul) F. Muell., Fragm. (Mueller) 6(47): 193 (1868).
*Paratrophis heterophylla* Blume, Mus. Bot. 2(1-8): 81. (1856).
8. ***Paratrophis pendulina* (Endl.) E.M. Gardner, comb. nov.** — based on *Morus pendulina* Endl., Prodromus Florae Norfolkicae 40 (1833) – *Pseudomorus pendulina* (Endl.) Stearn, J. Arnold Arbor. 28: 427 (1947) – *Pseudomorus brunoniana* var. *pendulina* (Endl.) Bureau, Ann. Sci. Nat. Bot. sér. 5, 11: 372 (1869) – *Streblus pendulinus* (Endl.) F. Muell., Fragmenta Phytographiae Australiae (1868).

*Morus brunoniana* Endl., Atakta Bot. t. 32 (1835) – *Streblus brunonianus* (Endl.) F. Muell., Fragm. Phyt. Australiae 6: 192 (1868) – *Pseudomorus brunoniana* (Endl.) Bureau, Ann. Sci. Nat., Bot. sér. 5, 11: 372 (1869)
*Pseudomorus brunoniana* var. *australiana* Bureau, Ann. Sci. Nat., Bot. sér. 5, 11: 373 (1869) – *Pseudomorus pendulina* var. *australiana* (Bureau) Stearn, J. Arnold Arbor. 28: 427 (1947).
*Pseudomorus brunoniana* var. *australiana* subvar. *castaneaefolia* Bureau, Ann. Sci. Nat., Bot. sér. 5, 11: 372 (1869).
*Pseudomorus brunoniana* var. *obtusa* Bureau, Ann. Sci. Nat., Bot. sér. 5, 11: 373 (1869) – *Pseudomorus pendulina var. obtusa* (Bureau) Stearn, J. Arnold Arbor. 28: 428 (1947), as var. *obtusata*.
*Pseudomorus sandwicensis* O. Deg., Fl. Hawaiiensis fam. 38 (1938) – *Pseudomorus brunoniana* var. *sandwicensis* (O. Deg.) Skottsb., Acta Horti Gothob. 15: 347 (1944) *– Pseuomorus pendulina* var. *sandwicensis* (O. Degener) Stearn, J. Arnold Arbor. 28: 428 (1947) – *Streblus sandwicensis* (O. Deg.) H. St. John, Pacific Trop. Bot. Gard. Mem. 1: 374. 1973. Note: This complex requires further investigation. Although the types of *Morus pendulina*, *Morus brunoniana*, *Pseudomorus sandwicensis*, and *Pseudomorus brunoniana* var. *obtusa* differ from one another, it is difficult to discern clear geographic patterns that warrant maintaining separate species, and we therefore provisionally treat them all as one widespread and variable species.
9. ***Paratrophis philippinensis*** (Bureau) Fern.-Vill., Nov. App. 98 (1880) — *Uromorus philippinensis* Bureau in A.DC., Prodr. 17: 237 (1873) — *Trophis philippinensis* (Bureau)

Corner, Gard. Bull. Singapore 19: 231 (1962).
*Sloetia minahassae* Koord., Versl. Minahasa 645 (1898).
*Paratrophis grandifolia* Elmer, Leafl. Philipp. Bot. 5: 1814 (1913).
*Calpidochlamys branderhorstii* Diels, Bot. Jahrb. Syst. 67: 173 (1935) — *Trophis*
*branderhorstii* (Diels) Corner, Gard. Bull. Singapore 19: 231 (1962).
*Calpidochlamys drupacea* Diels, Bot. Jahrb. Syst. 67: 173 (1935) — *Trophis drupacea*
(Diels) Corner, Gard. Bull. Singapore 19: 231 (1962). Note: This species is the only *Paratrophis* with fused tepals and one of only two whose tepals develop into a fleshy accessory fruit. The staminate inflorescences, however, are typical of the genus.
10. ***Paratrophis sclerophylla* (Corner) E.M. Gardner, comb. nov.** — based on *Streblus sclerophyllus* Corner, Blumea 18: 399 (1970).
11. ***Paratrophis smithii*** Cheeseman T.N.Z.L 20:148 (1888) – *Streblus smithii* (Cheeseman) Corner Gard. Bull. Singapore 19: 224 (1962). Note: Morphologically very close to *P. anthropophagorum*.
12. ***Paratrophis solomonensis* (Corner) E.M. Gardner, comb. nov.** — based on *Streblus solomonensis* Corner, Gard. Bull. Singapore 19: 224 (1962). Note:— Morphologically very close to *P. anthropophagorum*.

#### Sorocea

*Sorocea* A. St-Hil., Mém. Mus. Hist. Nat. 7: 473 (1821).

*Balanostreblus* Kurz, J. Asiat. Soc. Bengal., Pt. 2, Nat. Hist., 42: 247 (1873), p.p.
*Pseudosorocea* Baill., Hist. Pl. 6: 210 (1875).
*Trophisomia* Rojas Acosta, Bull. Acad. Inst. Geogr. Bot. 24: 211 (1914).
*Paraclarisia* Ducke, Arq. Serv. Florest. 1(1): 2 (1939).

Dioecious trees. ***Leaves*** alternate, distichous, lamina pinnately veined, 3-lobed to entire. ***Stipules*** free, lateral. ***Inflorescences*** solitary or paired in the leaf axils or below the leaves, racemose to spicate to subcapitate or uniflorous, bracts basally attached to peltate. Staminate flowers 4-lobed to 4-parted, tepals decussate-imbricate, stamens (3–)4, straight in bud, pistillode usually absent. Pistillate flowers tubular, 4-lobed to sub-entire, ovary basally adnate to the perianth, stigmas 2, short, usually tongue-shaped. ***Infructescences*** with drupaceous fruits enclosed by fleshy enlarged perianths, the latter red to orange, turning black at maturity. ***Seeds*** large, testa thin, embryo green, cotyledons very unequal, the smaller one minute and enclosed by the larger.

Species and distribution: 19 species in Central and South America from Mexico to Argentina.

Note: This species list follows the *Flora Neotropica* monograph (Berg 2001) with the addition of four new species and one status change published later. Names synonymized by Berg (2001) but recognized by Burger et al. (1962) are noted in parentheses. Complete synonymies can be found in those treatments.

*Sorocea* subg. *Sorocea*

***Sorocea affinis*** Hemsl., Biol. Cent.-Amer., Bot. 3: 150 (1883).
***Sorocea angustifolia*** Al.Santos & Romaniuc, Novon 24: 199 (2015).
***Sorocea bonplandii*** (Baill.) W.C. Burger, Lanj. & de Boer, Acta Bot. Neerl. 11: 465 (1962).
***Sorocea briquetii*** J.F.Macbr., Publ. Field Columb. Mus., Bot. Ser. 11: 16 (1931) (incl. *S. pileate* W.C. Burger).
***Sorocea carautana*** M.D.M.Vianna, Carrijo & Romaniuc, Novon 19: 549 (2009).
***Sorocea ganevii*** R.M.Castro, Neodiversity 1: 18 (2006).
***Sorocea guilleminiana*** Gaudich., Voy. Bonite, Bot. 3: t. 74 (1843) (incl. *S. klotzschiana* Baill. and *S. macrogyna* Lanj. & Wess. Boer).
***Sorocea hilarii*** Gaudich., Voy. Bonite, Bot. 3: t. 71 (1843) (incl. *S. racemosa* Gaudich.).
***Sorocea jaramilloi*** C.C.Berg, Novon 6: 241 (1996).
***Sorocea longipedicellata*** A.F.P. Machado, M.D.M. Vianna & Romaniuc, Syst. Bot. 38: 687 (2013).
***Sorocea muriculata*** Miq., C.F.P. von Martius & auct. suc. (eds.), Fl. Bras. 4(1): 113 (1853).
  **subsp. *muriculata*** (incl. *S. amazonica* Miq.)
  **subsp. *uaupensis*** (Baill.) C.C. Berg, Proc. Kon. Ned. Akad. Wetensch. C 88: 387 (1985) (incl. *S. guayanensis* W.C. Burger).
***Sorocea pubivena*** Hemsl., Biol. Cent.-Amer., Bot. 3: 150 (1883).
  **subsp. *pubivena*** (incl. *S. cufodontii* W.C. Burger)
  **subsp. *hirtella*** (Mildbr.) C.C. Berg, Novon 6: 243 (1996) (incl. *S. opima* J.F. Macbr.)
  **subsp. *oligotricha*** (Akkermans & C.C. Berg) C.C. Berg, Novon 6: 243 (1996).
***Sorocea ruminata*** C.C.Berg, Novon 6: 244 (1996).
***Sorocea sarcocarpa*** Lanj. & Wess. Boer, Acta Bot. Neerl. 11: 452 (1962).
***Sorocea steinbachii*** C.C. Berg, Proc. Kon. Ned. Akad. Wetensch. C 88: 385 (1985).
***Sorocea trophoides*** W.C. Burger, Acta Bot. Neerl. 11: 450 (1962).
*Sorocea* subg. *Paraclarisia* (Ducke) W.C. Burger, Lanj. & Wess. Boer, Acta Bot. Neerl. 11: 468 (1962).

***Sorocea duckei*** W.C.Burger, Acta Bot. Neerl. 11: 473 (1962).
***Sorocea sprucei*** (Baill.) J.F.Macbr., Publ. Field Mus. Nat. Hist., Bot. Ser. 11: 16 (1931).
  **subsp. *sprucei*** (incl. *S. arnoldoi* Lanj. & Wess. Boer).
  **subsp. *saxicola*** (Hassl.) C.C.Berg, Proc. Kon. Ned. Akad. Wetensch., Ser. C., Biol. Med. Sci. 88: 391 (1985).
***Sorocea subumbellata*** (C.C. Berg) Cornejo, Novon 19: 297 (2009).

#### Taxotrophis

*Taxotrophis* Blume, Ann. Mus. Bot. Lugduno-Batavi 2 77 (1856); Hutch., Bull. Misc. Inform. Kew 147 (1918).

*Streblus* Lour. sect. *Taxotrophis* (Blume) Corner, Gard. Bull. Singapore 19: 218 (1962); Berg et al., Fl. Males. Ser. 1, Vol. 17, Pt. 2 (2006).

Monoecious or dioecious trees or shrubs, usually with lateral or terminal thorns. ***Leaves*** distichous, pinnately veined, petioles adaxially pubescent. Stipules free, lateral. ***Inflorescences*** axillary, solitary or paired, with an abaxial sterile groove, interfloral bracts basally attached, flowers with 4 free tepals, valvate. Staminate inflorescences spicate to sub-capitate, flowers with filaments inflexed in bud, pistillode present. Pistillate inflorescences uniflorous or sub-spicate, flowers usually pedicellate, stigma bifid, arms equal. ***Fruits*** drupaceous, up to ca. 1 cm long, usually loosely enclosed by the enlarged persistent tepals. ***Seeds*** up to ca. 8 × 6 mm, cotyledons subequal to unequal.

Species and distribution: Six species, ranging from Sri Lanka to New Guinea.

1. ***Taxotrophis ilicifolia*** (Kurz) S.Vidal, Revis. Pl. Vasc. Filip. 249 (1886) – *Balanostreblus ilicifolia* Kurz, J. Asiat. Soc. Bengal, Pt. 2, Nat. Hist. 42(4): 248

(1874) – *Streblus ilicifolius* (Kurz) Corner, Gard. Bull. Singapore 19: 227 (1962). *Pseudotrophis laxiflora* Warb., Bot. Jahrb. Syst. 13 294 (1891).
*Taxotrophis obtusa* Elmer, Leafl. Philipp. Bot. 5 1813 (1913).
*Taxotrophis laxiflora* Hutch., Bull. Misc. Inform. Kew 151 (1918). – *Streblus laxiflorus* (Hutch.) Corner, Gard. Bull. Singapore 19 229 (1962).
*Taxotrophis triapiculata* Gamble, Bull. Misc. Inform. Kew 188 (1913).
*Taxotrophis eberhardtii* Gagnep., Fl. Indo-Chine 5 700 (1928).
*Taxotrophis macrophylla* auct. non Boerl.: Burkill, Dict. Econ. Prod. Malay Penins. 2126 (1935).
2. ***Taxotrophis macrophylla*** (Blume) Boerl., Handl. Fl. Ned. Ind. 3: 359 (1900).

*Streblus macrophyllus* Blume, Ann. Mus. Bot. Lugduno-Batavi 2: 80 (1856) — *Diplocos ? macrophyllus* (Blume) Bureau in A.DC., Prodr. 17: 216 (1873).
*Pseudotrophis mindanaensis* Warb. in Perkins, Fragm. Fl. Philipp. 1: 165 (1905); Elmer, Leafl. Philipp. Bot. 5: 1815 (1913), ‘*Taxatrophis mindanaensis*’ in nota.
*Paratrophis caudata* Merr., Philipp. J. Sci., 1, Suppl. 183 (1906).
*Taxotrophis balansae* Hutch., Bull. Misc. Inform. Kew 151 (1918).
*Dimerocarpus brenieri* Gagnep., Bull. Mus. Hist. Nat. (Paris) 27: 441 (1921).
3. ***Taxotrophis perakensis* (Corner) E.M. Gardner, comb. nov.**, based on *Streblus perakensis* Corner, Gard. Bull. Singapore 19: 223 (1962). Note: Although Corner (1962) considered *S. perakensis* species part of section *Paratrophis*, albeit with some hesitation, Berg et al. (2006), whom we follow, placed it in *Taxotrophis* based on the spines that appear on some specimens. This is a rather variable species that requires further investigation to properly elucidate its limits and affinities.
4. ***Taxotrophis spinosa*** Steenis in Backer & Bakh.f., Fl. Java 2 (1965) 16.

*Urtica spinosa* Blume, Bijdr. (1825) 507. — *Streblus spinosus* (Blume) Corner, Gard. Bull. Singapore 19: 229 (1962).
*Taxotrophis javanica* Blume, Ann. Mus. Bot. Lugduno-Batavi 2: 77, t. 26 (1856).
5. ***Taxotrophis taxoides*** (B. Heyne ex Roth) W.L. Chew ex E.M. Gardner, comb. nov.

*Trophis taxoides* B. Heyne ex Roth, Nov. Pl. Sp. 368 (1821). — *Trophis taxiformis* Spreng., Syst. Veg. 3 902 (1826), nom. nov. illeg. — *Streblus taxoides* (B. Heyne ex Roth) Kurz, Forest Fl. Burma 2: 465 (1877) — *Phyllochlamys taxoides* (B. Heyne ex Roth) Koord., Exkurs.-Fl. Java 2: 89 (1912).
*Trophis spinosa* Roxb., Fl. Ind., ed. Carey 3: 762 (1832), non Willd. 1806, nec Blume 1826. — *Epicarpurus spinosus* (Roxb.) Wight, Icon. Pl. Ind. Orient. 6: 7, t. 1962 (1853), p.p. — *Phyllochlamys spinosa* (Roxb.) Bureau in A.DC., Prodr. 17: 218 (1873);
*Epicarpurus timorensis* Decne., Nouv. Ann. Mus. Hist. Nat. 3: 499, t. 21 (1834).
*Taxotrophis roxburghii* Blume, Ann. Mus. Bot. Lugduno-Batavi 2: 78 (1856)
*Streblus microphyllus* Kurz, Prelim. Rep. Forest Pegu App. A, cxviii; App. B, 84 (1875); — *Streblus taxoides* (B. Heyne ex Roth) Kurz var. *microphylla* (Kurz) Kurz, Forest Fl. Burma 2: 465 (1877).
*Phyllochlamys wallichii* King ex Hook.f., Fl. Brit. India 5: 489 (1888);
*Phyllochlamys taxoides* (B. Heyne ex Roth) Koord. var. *parvifolia* Merr., Enum. Philipp. Flow. Pl. 2: 38 (1923).
*Taxotrophis poilanei* Gagnep., Fl. Indo-Chine 5: 701 (1928).
*Taxotrophis crenata* Gagnep., Fl. Indo-chine 5: 702, t. 82 (1928). — *Streblus crenatus*
(Gagnep.) Corner, Gard. Bull. Singapore 19: 226 (1962).
*Phyllochlamys tridentata* Gagnep., Fl. Indo-Chine 5: 714 (1928). Note: In the 1950s, Dr. Chew Wee Lek annotated quite a lot of specimens at K and SING with the new combination *Taxotrophis taxoides*. However, the combination was never published, probably because the need for the combination was obviated in 1962 when Corner, his doctoral supervisor, reduced *Taxotrophis* to a section of *Streblus*.
6. ***Taxotrophis zeylanica*** (Thwaites) Thwaites, Enum. Pl. Zeyl. [Thwaites] 264 (1861).

*Epicarpurus zeylanicus* Thwaites, Hooker’s J. Bot. Kew Gard. Misc. 4: 1 (1852) — *Diplocos zeylanica* (Thwaites) Bureau, Prodr. [A. P. de Candolle] 17: 215 (1873)
— *Streblus zeylanicus* (Thwaites) Kurz, Forest Fl. Burma 2: 464 (1877).
*Taxotrophis caudata* Hutchinson, Bull. Misc. Inform. Kew 1918(4): 149 (1918).

#### Trophis

*Trophis* Lour., P. Browne, Civ. Nat. Hist. Jamaica 357 (1756), nom. cons.

*Bucephalon* L. Sp. Pl. 1190 (1753), nom. rejic.

*Skutchia* Pax & K. Hoffm. ex C.V. Morton, J. Wash. Acad. Sci. 27: 306 (1937).

Dioecious trees or shrubs. ***Leaves*** distichous, pinnately veined. Stipules free, lateral. ***Inflorescences*** axillary or just below the leaves, solitary or paired, interfloral bracts basally attached. Staminate inflorescences spicate to racemose with an abaxial sterile groove, tepals 4, basally connate, stamens 4, filaments inflexed in bud, pistillode present. Pistillate inflorescences spicate to racemose or subcapitate, tepals 4, connate, forming a tubular perianth, ovary adnate to the perianth or not, stigma bifid, arms equal. ***Fruits*** drupaceous, up to ca. 1.5 cm long, adnate to the perianth or not, the perianth enlarged and fleshy or not. ***Seeds*** up to ca. 1 cm long, cotyledons equal.

Species and distribution: Five species in the neotropics.

*Trophis* P. Browne sect. *Trophis*

1. ***Trophis cuspidata*** Lundell, Amer. Midl. Naturalist 19: 427 (1938).
2. ***Trophis mexicana*** (Liebm.) Bureau, Prodr. [A. P. de Candolle] 17: 253 (1873).
3. ***Trophis noraminervae*** Cuevas & Carvajal, Acta Bot. Mex. 47: 2 (1-7, fig.) (1999).
4. ***Trophis racemosa*** Urb., Symb. Antill. (Urban). 4(2): 195 (1905).

*Trophis* P. Browne sect. *Echinocarpa* C.C. Berg, Proc. Kon. Ned. Akad. Wetensch., Ser. C, Biol. Med. Sci. 91: 353 (1988).

5 ***Trophis involucrata*** W.C. Burger, Phytologia 26: 432 (1973).

#### Tribe OLMEDIEAE

Olmedieae Trécul, Ann. Sci. Nat., Bot. sér. 3, 8 (1847) 126

Castilleae C.C. Berg, Acta Bot. Neerl. 26 (1977) 78

Strebleae Bureau in DC., Prodr. 17: 215 (1873), p.p.

Tree or shrubs, monoecious or dioecious (or androdioecious), mostly with self-pruning branches. ***Leaves*** alternate or opposite, distichous or spirally arranged; stipules lateral to amplexicaul. ***Inflorescences*** mostly unisexual, capitate, mostly discoid to urceolate, involucrate, tepals mostly 4, connate or not. Staminate inflorescences usually many-flowered; stamens 4 or fewer, with filaments straight or less often inflexed in bud, pistillode mostly absent. Pistillate inflorescences one to many-flowered, ovary free or not, stigmas 2, filiform. ***Fruits*** mostly drupaceous, mostly enclosed by a fleshy perianth or embedded in a fleshy receptacle. ***Seeds*** with or without endosperm, testa thin, vascularized, cotyledons mostly equal.

Genera and distribution: 13 genera with 63 species. Eight neotropical genera: *Castilla* (3 spp.), *Helicostylis* (7), *Maquira* (4), *Naucleopsis* (22), *Olmedia* (1), *Perebea* (9), *Poulsenia* (1), and *Pseudolmedia* (9); and five Paleotropical genera: the widespread *Antiaris* (1); *Antiaropsis* (2) in New Guinea; *Mesogyne* (1) in Africa; *Sparattosyce* (1) in New Caledonia; and *Streblus* (3) in South to Southeast Asia.

#### Olmedia

*Olmedia* Ruiz & Pav., Syst. Veg. Fl. Peruv. Chil.1:257.1798. Type — *Olmedia aspera* Ruiz & Pav.

*Trophis* section *Olmedia* (Ruiz & Pav.) Berg, Proc. Kon. Ned. Acad. Wetensch. C. 91: 354. 1988.

Dioecious trees or shrubs. ***Leaves*** distichous, lamina pinnately veined. ***Stipules*** free, not fully amplexicaul. ***Inflorescences*** unisexual, with a well-developed involucre. Staminate inflorescences discoid, multiflorous; tepals 4, valvate, stamens 4, inflexed in bud, pistillode absent. Pistillate inflorescences usually uniflorous, perianth tubular, 4-dentate, ovary free, stigmas 2, equal. ***Fruits*** drupaceous, surrounded by fleshy persistent perianth and subtended by spreading, fleshy involucral bracts. ***Seeds*** ca. 5 mm long, cotyledons equal.

Species and distribution: One species, in the Neotropics.

1. ***Olmedia aspera*** Ruiz & Pav., Syst. Veg. Fl. Peruv. Chil.1:257.1798

*Olmedia caucana* Pittier, Contr. U.S. Natl. Herb. 13:434. 1912. *— Trophis caucana* (Pitttier) C.C. Berg, Proc. Kon. Ned. Acad. Wetensch. C. 91: 354. 1988
*Olmedia poeppigiana* Klotzsch, Linnaea 20: 525. 1847, as a synonym of *O. aspera* Poeppig & Endlicher, Nov. Gen. 2: 31. 1838, based on Poeppig s.n. or 1267, non *O.poeppigiana* Martius Flora (or Bot. Zeit) 24 (Beibl. 2): 93. 1841 (= *Helicostylis tomentosa* (Poeppig & Endlicher) Rusby).
*Olmedia falcifolia* Pittier, Contr. U.S. Natl. Herb. 13: 435. 1912.
*Trophis aurantiaca* Herzog, Repert. Spec. Nov. 7: 51. 1909

#### Streblus

*Streblus* Lour., Fl. Cochinch. (1790) 615.

*Achymus* Juss., Dict. Sci. Nat. 1, Suppl. 31 (1816).

*Epicarpurus* Blume, Bijdr. 488 (1825).

*Albrandia* Gaudich. in Freyc., Voy. Uranie, Bot. 509 (1830).

*Calius* Blanco, Fl. Filip. 698 (1837).

*Teonongia* Stapf, Hooker’s Icon. Pl. 30: t. 2947 (1911).

*Diplothorax* Gagnep., Bull. Soc. Bot. France 75 98 (1928).

Trees or shrubs, dioecious or monoecious. ***Leaves*** distichous, lamina pinnately veined. ***Stipules*** free, lateral. ***Inflorescences*** bisexual or unisexual, capitate, with a rudimentary involucre, bracts basally attached. Staminate inflorescences discoid capitate, multiflorous; tepals 4, imbricate, stamens 4, inflexed in bud, pistillode present but small. Pistillate inflorescences usually uniflorous, tepals 4, ovary free, stigmas 2, equal. ***Fruits*** drupaceous, up to ca. 8 mm long, initially enclosed by enlarged but not fleshy tepals, which may open later. ***Seeds*** ca. 5 mm long, cotyledons equal or very unequal.

Species and distribution: three species from India to South China and from mainland Southeast Asia to the Philippines and the Moluccas.

1. ***Streblus asper*** (Retz.) Lour., Fl. Cochinch. 2: 615 (1790).

*Trophis aspera* Retz., Observ. Bot. 5 (1788) – *Epicarpurus asper* (Retz.) Steud. Nomencl. Bot. ed. 2, 1: 556 (1840).
*Trophis cochinchinensis* Poir., Encycl. 8 (1808) 123.
*Epicarpurus orientalis* Blume, Bijdr. (1825) 488
*Calius lactescens* Blanco, Fl. Filip. (1837) 698; ed. 3, 3 (1879) 1103, t. 171. — *Streblus lactescens* (Blanco) Blume, Ann. Mus. Bot. Lugduno-Batavi 2 (1856) 80.
*Achymus pallens* Sol. ex Blume, Ann. Mus. Bot. Lugduno-Batavi 2 (1856) 79.
*Cudrania crenata* C.H. Wright, J. Linn. Soc., Bot. 26 (1899) 469. — *Vanieria crenata* (C.H. Wright) Chun, J. Arnold Arbor. 8 (1927) 21.
*Diplothorax tonkinensis* Gagnep., Bull. Soc. Bot. France 75 (1928) 98.
2. Note: Retzius’s *Trophis aspera* predated Loureiro’s *Streblus asper* by two years, making the latter an implied new combination under Article 41.1 of the Code (cf. Ex. 10). Synonymies of the *S. asper* have sometimes erroneously included *Trophis aculeata* Roth, which is actually *Maclura spinosa* (Roxb. ex Willd.) C.C. Berg.
3. ***Streblus celebensis*** C.C. Berg, Blumea 50 547 (2005).
4. ***Streblus tonkinensis*** (Eberh. & Dubard) Corner, Gard. Bull. Singapore 19 (1962): 228. *Bleeokrodea tonkinensis* Eberh. & Dubard, Compt. Rend. Hebd. SÈances Acad. Sci. 145 (1907): 632. — *Teonongia tonkinensis* (Eberh. & Dubard) Stapf.

#### Tribe DORSTENIEAE

Dorstenieae Gaudich. in Freyc., Voy. Uranie, Bot. (1830).

Broussonetieae Gaudich. in Freyc., Voy. Uranie, Bot. 508 (1830).

Brosimeae Trécul, Ann. Sci. Nat., Bot. sér. 3, 8: 146 (1847).

Fatoueae Engl., Nat. Pflanzenfam. 3, 1: 71 (1888).

Tree, shrubs, lianas, and herbs, monoecious or less often dioecious. ***Leaves*** alternate or less commonly (sub)opposite, distichous or spirally arranged; stipules lateral to fully amplexicaul. ***Inflorescences*** unisexual or bisexual, cymose, spicate, globose, or discoid to turbinate or cup-shaped, multiflorous or uniflorous (the latter in pistillate inflorescences only), bracteate or not, interfloral bracts mostly peltate. Staminate flowers with tepals (1–)2–4 or absent, stamens 1–4 with filaments straight or inflexed in bud, pistillode present or (more often) absent. Pistillate flowers free or connate or embedded in the receptacle, tepals 2–4, ovary free or not, stigmas 1 or 1, equal or unequal. ***Fruits*** drupaceous or drupe-like due to a persistent fleshy perianth and/or receptacle, the whole endocarp unit often ballistically ejected from the infructescence. ***Seeds*** with or without endosperm, large or small; cotyledons equal or unequal.

Genera and distribution: eleven genera and 63 species with a worldwide distribution: *Bleekrodea* (Africa and Southeast Asia)*, Bosqueiopsis* (Africa)*, Brosimum* (Neotropics)*, Broussonetia* (Asia to Oceania; introduced worldwide)*, Dorstenia* (Africa and South America, with one species in India)*, Fatoua* (Madagascar and Japan to New Caledonia; introduced worldwide), *Helianthostylis* (Neotropics)*, Malaisia* (Southeast Asia to New Caledonia)*, Scyphosyce* (Africa)*, Sloetia* (Southeast Asia)*, Sloetiopsis* (Africa), *Trilepisium* (Africa), *Trymatococcus* (Neotropics), *Utsetela* (Africa).

#### Sloetiopsis

*Sloetiopsis* Engl., Bot. Jahrb. 39: 573 (1907).

*Neosloetiopsis* Engl., Bot. Jahrb. 51: 426 (1914)

*Streblus* auct. non Lour., C.C. Berg, Proc. Kon. Ned. Acad. Wetensch. C. 91: 357. 1988.

Tree or shrubs, dioecious (or monoecious). ***Leaves*** distichous, pinnately veined, cystoliths present; stipules free, nearly amplexicaul. ***Inflorescences*** unisexual (or bisexual); staminate inflorescences spicate with an abaxial sterile groove, bracts mostly peltate, tepals 4, free or basally connate filaments inflexed in bud, pistillode small; pistillate inflorescences uniflorous, bracts mostly basally attached, tepals 4, free, imbricate, 2 stigmas. ***Fruits*** drupaceous, fleshy endocarp dehiscent, tepals enlarged and persistent but not fleshy. ***Seeds*** globose, ca. 4 mm, endocarp coriaceous with a hard disc against the hilum, testa vascularized with a thick apical cap, cotyledons equal.

1. ***Sloetiopsis usambarensis*** Engl., Bot. Jahrb. 39: 573 (1907) — *Streblus usambarensis* (Engl.) C.C. Berg, Proc. Kon. Ned. Acad. Wetensch. C. 91: 357. 1988.

*Neosloetiopsis kamerunensis* Engl., Bot. Jahrb. 51: 426 (1914).

Note. The nearly amplexicaul stipules and occasional bisexual inflorescences reflect an affinity with the Southeast Asian *Sloetia*, also a member of the Dorstenieae.

#### Tribe PARARTOCARPEAE

Parartocarpeae N.J.C. Zerega & E.M. Gardner Zerega, Phytotaxa 388, 253–265 (2019). Shrubs to large trees, monoecious or dioecious; abundant white exudate. ***Leaves*** distichous or spirally arranged; simple; entire; pinnately veined; thin to thick coriaceous; glabrous, pubescent, scabrid, or hispid pubescent. ***Stipules*** axillary, simple or paired, lateral. ***Inflorescences*** solitary or paired in leaf axils, unisexual (or bisexual with a single apical pistillate flower), uniflorous, cymose, or capitate with stamens or ovaries sunken into the receptacle; pedunculate; involucre of 3–8 triangular bracts, basally connate. Staminate inflorescences with numerous flowers, tepals 4–5, stamens 5 and normally positioned or 1– 3 in cavities in the receptacle with the anthers exerted through perforations in the upper surface of the receptacle, filaments free or united. Pistillate inflorescences uniflorous or (sub)globose with ovaries solitary in each cavity, unilocular, the style apical with a short exerted stigma. ***Fruits*** drupaceous, enclosed by persistant tepals, or aggregated into syncarps formed by the enlargement of the entire inflorescence.

Genera and distribution: Three genera and five species, from Southern China to the Solomon Islands: *Hullettia*, *Parartocarpus*, and *Pseudostreblus*.

#### Pseudostreblus

*Pseudostreblus* Bureau, in A.P. de Candolle, Prodr. 17: 220 (1873).

*Streblus* Lour. sect. *Pseudostreblus* (Bureau) Corner, Gard. Bull. Singapore 19: 217 (1962).

Monoecious trees. ***Leaves*** distichous, pinnately veined. Stipules free, lateral. ***Inflorescences*** axillary. Staminate inflorescences cymose, tepals 5, imbricate, filaments inflexed in bud but apparently straightening gradually upon anthesis, pistillode minute, conical, pubescent, interfloral bracts few, basally attached. Pistillate inflorescences uniflorous, pedunculate, involucral bracts 3, ±connate, tepals 4, imbricate, stigma bifid, arms equal. ***Fruits*** drupaceous, ca. 1 cm long, with tepals enlarged and loosely enclosing the fruit. ***Seeds*** ellipsoid, ca. 6 × 8 mm, cotyledons unequal.

Species and distribution: One, in South to Southeast Asia and Southern China.

Note: *Pseudostreblus* is remarkable for its five-parted staminate flowers.

1. ***Pseudostreblus indicus* Bureau**, in A.P. de Candolle, Prodr. 17: 220 (1873).

*Streblus indicus* (Bureau) Corner, Gard. Bull. Singapore 19: 226 (1962).

## Conclusion

The revisions presented here, based on the best phylogenetic evidence available to date, provide for seven monophyletic tribes of Moraceae and ten monophyletic genera within Moreae. We hope that the resulting taxonomic and phylogenetic framework will provide a solid foundation for further research into the systematics and evolution of Moraceae.

## Author contributions

EMG led the writing of the manuscript, did field, lab, and herbarium work, conducted all phylogenetic analyses, and was responsible for the taxonomic treatment. MG performed lab work for the Moraceae333 samples. RC, SD, NE, and OM performed lab work for the PAFTOL samples, and OM curated the PAFTOL data. DA collaborated on all herbarium work at BO. Sahromi facilitated access and participated in field collections at the Bogor Botanical Gardens. NJCZ provided overall guidance on Moraceae and facilitated sequencing at Northwestern University. WJB and FF acquired funding for and supervised the PAFTOL project. AM performed herbarium sampling at K and supervised the Moraceae portion of the PAFTOL project. EMG designed the overall study with input from ALH and NJCZ. ALH supervised the overall study. All authors commented on and contributed to the manuscript.

## Acknowledgements

This work was supported by the United States National Science Foundation (DBI award number 1711391, and DEB award numbers 0919119 and 1501373); the SYNTHESYS Project http://www.synthesys.info/ which is financed by European Community Research Infrastructure Action under the FP7 “Capacities” Program”; the National Parks Board of Singapore through a *Flora of Singapore* Fellowship and a Research Fellowship; grants from the Calleva Foundation and the Sackler Trust to the Plant & Fungal Trees of Life Project at the Royal Botanic Gardens, Kew. We thank the Pritzker Laboratory for Molecular Systematics at the Field Museum of Natural History (K. Feldheim) for access to laboratory and sequencing facilities; S. Knapp and J. Wajer (BM) for helpful consultation on nomenclatural issues; W. Clement for helpful comments on the circumscription of the Olmedieae; the Bogor Botanical Gardens, Singapore Botanic Gardens, and The Morton Arboretum for access to living collections; and the following herbaria for access to collections for examination and/or sampling: BM, BKF, BO, CLM, F, IBSC, K, FTBG, L, MO, NY, SING, SAN, SAR, SNP, and U. Finally, we gratefully acknowledge the careful work of past scholars of Moraceae, in particular the late C.C. Berg and E.J.H. Corner.

