## Supplementary figures and images for "Repeated parallel losses of inflexed stamens in Moraceae: phylogenomics and generic revision of the tribe Moreae and the reinstatement of the tribe Olmedieae (Moraceae)"

### Figure S1

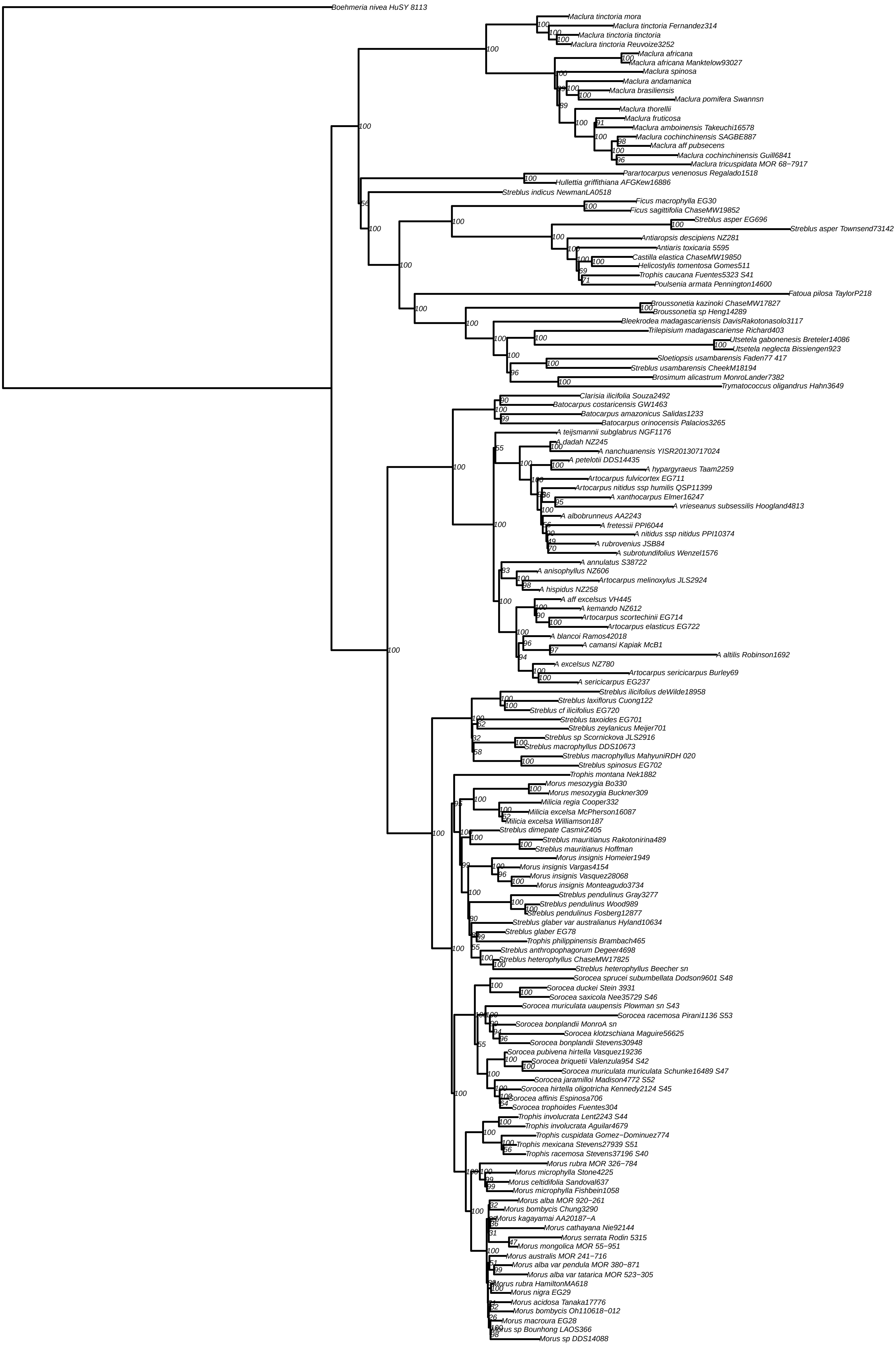

### Figure S2

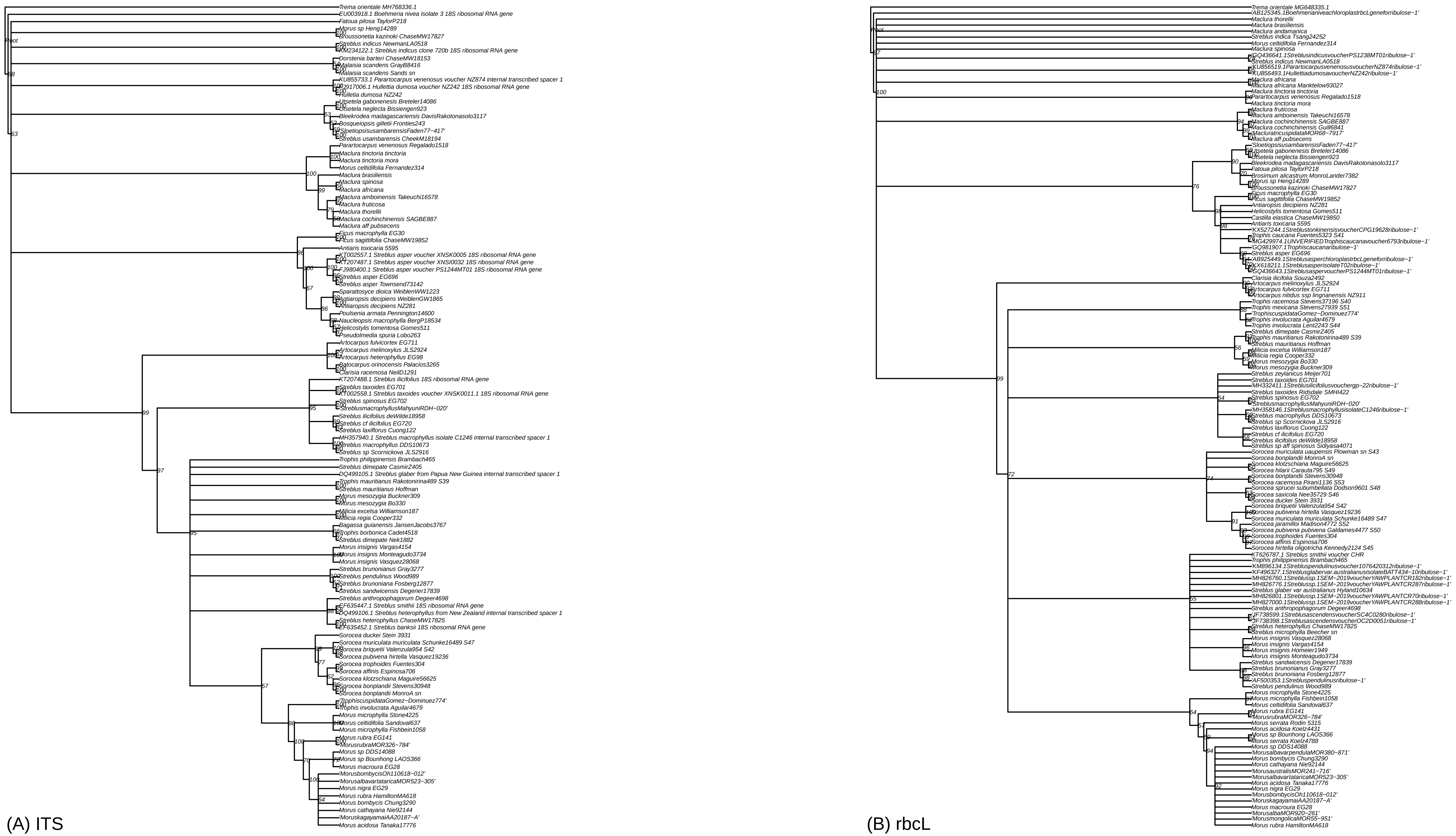

### Figure S3

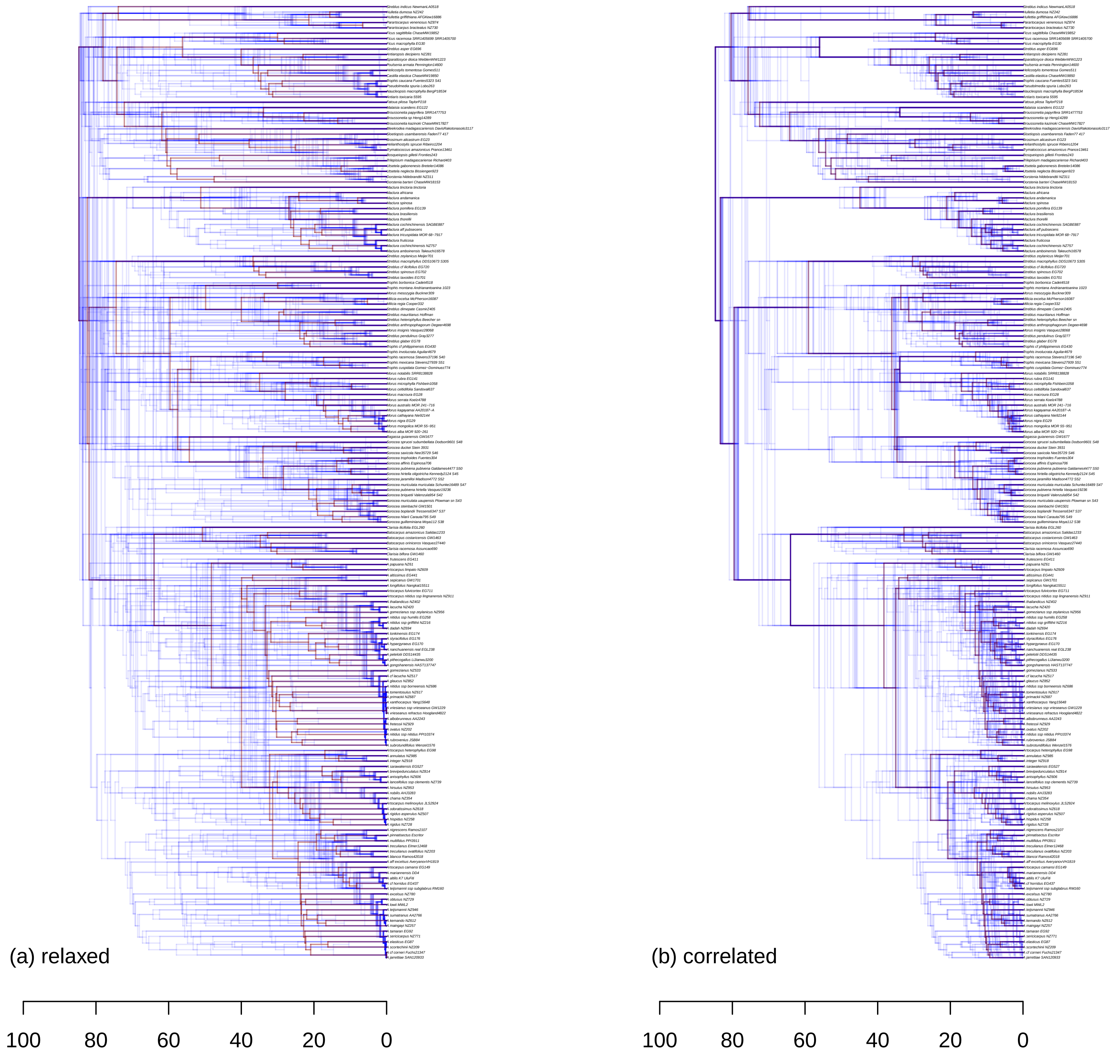

### Figure S4

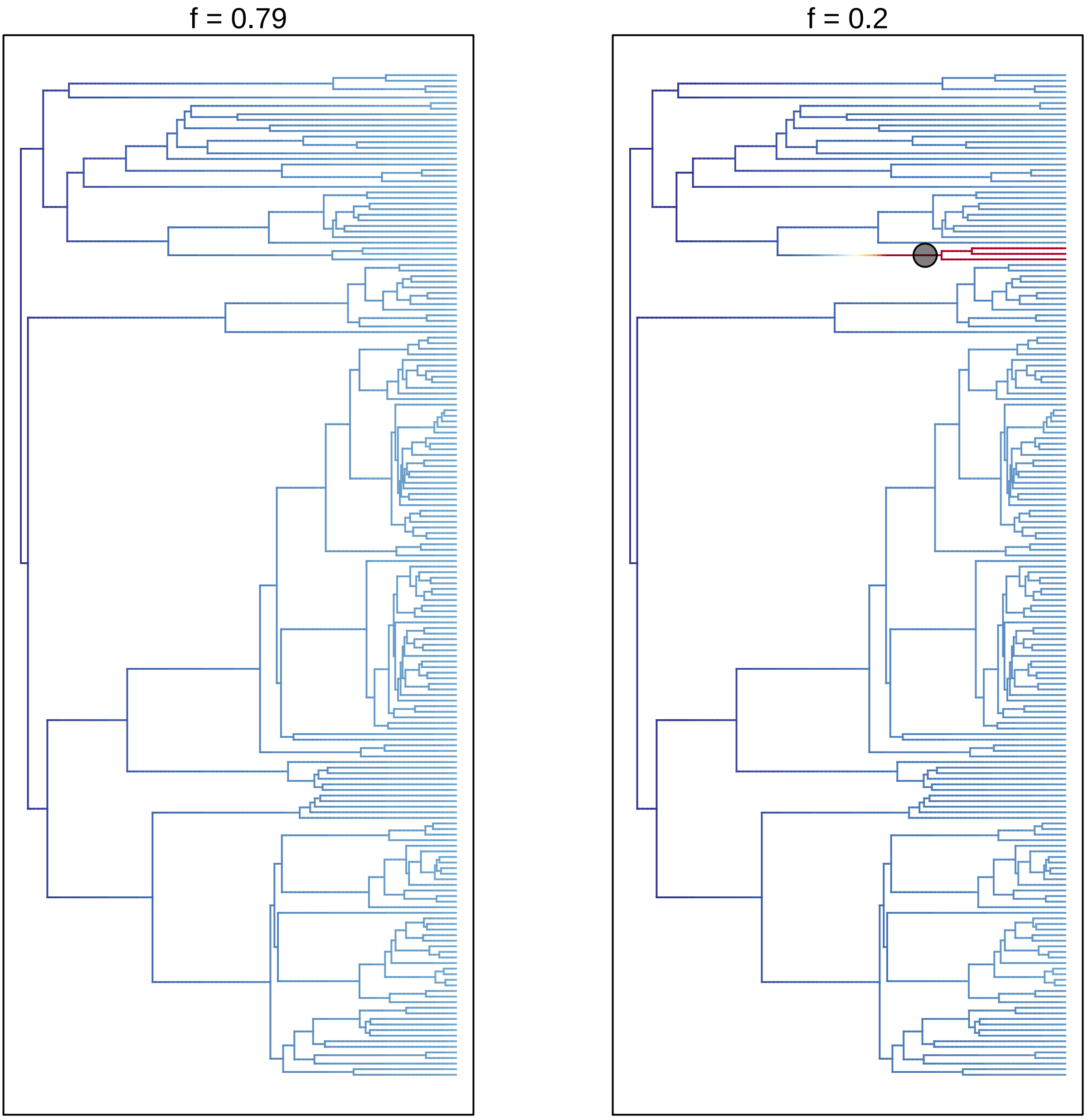
